# Activity-dependent subcellular compartmentalization of dendritic mitochondria structure in CA1 pyramidal neurons

**DOI:** 10.1101/2023.03.25.534233

**Authors:** Daniel M. Virga, Stevie Hamilton, Bertha Osei, Abigail Morgan, Emiliano Zamponi, Natalie J. Park, Victoria L. Hewitt, David Zhang, Kevin C. Gonzalez, Erik Bloss, Franck Polleux, Tommy L. Lewis

**Affiliations:** Department of Neuroscience, Columbia Medical School, New York, NY- USA; Mortimer B. Zuckerman Mind Brain Behavior Institute, Columbia University, New York, NY- USA; Aging & Metabolism Program, Oklahoma Medical Research Foundation, Oklahoma City, OK, USA; Neuroscience, Oklahoma University Health Science Campus, Oklahoma City, OK, USA; The Jackson Laboratory, 600 Main Street, Bar Harbor, ME 04609, USA

## Abstract

Neuronal mitochondria play important roles beyond ATP generation, including Ca^2+^ uptake, and therefore have instructive roles in synaptic function and neuronal response properties. Mitochondrial morphology differs significantly in the axon and dendrites of a given neuronal subtype, but in CA1 pyramidal neurons (PNs) of the hippocampus, mitochondria within the dendritic arbor also display a remarkable degree of subcellular, layer-specific compartmentalization. In the dendrites of these neurons, mitochondria morphology ranges from highly fused and elongated in the apical tuft, to more fragmented in the apical oblique and basal dendritic compartments, and thus occupy a smaller fraction of dendritic volume than in the apical tuft. However, the molecular mechanisms underlying this striking degree of subcellular compartmentalization of mitochondria morphology are unknown, precluding the assessment of its impact on neuronal function. Here, we demonstrate that this compartment-specific morphology of dendritic mitochondria requires activity-dependent, Camkk2- dependent activation of AMPK and its ability to phosphorylate two direct effectors: the pro-fission Drp1 receptor Mff and the recently identified anti-fusion, Opa1-inhibiting protein, Mtfr1l. Our study uncovers a new activity-dependent molecular mechanism underlying the extreme subcellular compartmentalization of mitochondrial morphology in dendrites of neurons *in vivo* through spatially precise regulation of mitochondria fission/fusion balance.

## INTRODUCTION

Neurons are among the largest, most complex and highly polarized cell types found in nature. The two main neuronal compartments, dendrites and the axon, differ dramatically in their overall architecture, morphology and functional properties. This striking degree of polarization requires differential targeting of mRNA and proteins, differences in organelle composition, such as specific classes of endosomes, as well as compartment-specific organelle morphology and dynamics ^1–5^. Two of the most abundant and complex organelles found in neurons, mitochondria and the endoplasmic reticulum, display a remarkable degree of morphological specialization between axonal and dendritic compartments ^5–10^: in the axon of long-projecting mammalian cortical pyramidal neurons (PNs), mitochondria are small (∼1 micron in length) and selectively localized to presynaptic boutons, whereas in the dendrites of the same neurons, mitochondria are long and tubular, filling a large fraction of the dendritic volume.

This high degree of polarization not only applies to these two broad neuronal compartments, but also to sub-compartments within axons and dendrites. Our recent work has identified a dramatic sub-cellular compartmentalization of mitochondrial morphology within the dendrites of CA1 PNs of the hippocampus *in vivo*^11^. Unlike other classes of PNs, such as cortical layer 2/3 PNs for example ^7^, CA1 PNs dendritic mitochondria are small and relatively uniform in shape in the dendritic compartments proximal to the soma (basal and apical oblique dendrites) but are significantly more tubular and occupy a larger fraction of the distal apical tuft compartment ^11^. Interestingly, this subcellular compartmentalization of mitochondrial morphology corresponds directly to the spatial segregation of presynaptic inputs received by CA1 PNs: basal dendrites in stratum oriens (SO) receive presynaptic inputs from CA2 and CA3 PNs, apical oblique dendrites in the stratum radiatum (SR) receive presynaptic inputs from CA3 PNs, while distal apical tuft dendrites in the stratum lacunosum moleculare (SLM) receive inputs from the entorhinal cortex (EC).

Mitochondrial morphology in dendrites of pyramidal neurons was previously shown to be regulated by synaptic activity such that neuronal depolarization increased dendritic mitochondria fission through Ca^2+^-dependent activation of the small-GTPase Dynamin-related protein 1 (Drp1) ^12–15^. However, most of these results were obtained *in vitro* and therefore their relevance for mitochondrial morphology of specific neuronal subtypes *in vivo* is largely unknown. In addition, Drp1 was recently shown to also be involved in regulating endocytosis in neurons ^16, 17^ obscuring the interpretation of these earlier results and leaving the molecular mechanisms whereby neuronal depolarization and/or synaptic activity could regulate dendritic mitochondrial morphology *in vivo* unresolved. Drp1 is a cytoplasmic protein recruited to the outer mitochondrial membrane (OMM) in order to promote constriction and fission in concert with so-called Drp1 ‘receptors’ present at the OMM including mitochondrial fission factor (Mff), fission 1 (Fis1), and mitochondrial dynamics proteins of 49 and 51 kDA (MiD49/MiD51) ^18–20^, with Mff being the most dominant and universal Drp1 receptor in mammalian cells. AMP-activated kinase (AMPK, also called Protein kinase AMP-activated or Prka) is a heterotrimer composed of a catalytic α subunit, an adaptor subunit β and the γ subunit that binds to AMP/ADP. In most cells, AMPK plays a central role as a metabolic sensor activated when ATP levels drop in cells (i.e., when AMP/ADP levels increase), and once activated AMPK phosphorylates many downstream substrates involved in restoring ATP levels and/or decreasing activity of pathways that consume significant levels of ATP such as protein synthesis ^21, 22^. In the context of metabolic stress, AMPK promotes mitochondrial fission by directly phosphorylating Mff^23^. Interestingly, in mammalian neurons AMPK is not only a metabolic sensor but also an activity-regulated kinase being phosphorylated and catalytically activated by Calcium/calmodulin dependent protein kinase kinase 2 (Camkk2), which is induced by increases in intracellular Ca^2+^ levels following either opening of voltage-gated Ca^2+^ channels (VGCC) induced by neuronal depolarization, or activation of N-methyl D-aspartate (NMDA) receptors ^11, 24, 25^. We have recently shown that Aβ42 oligomers trigger overactivation of Camkk2 leading to excessive activation of AMPK which phosphorylates Mff to trigger dendritic mitochondrial fission in apical tufts of CA1 PNs dendrites *in vivo* ^11^.

However, the physiological, developmental, and molecular mechanisms underlying this striking degree of mitochondrial compartmentalization in dendrites of CA1 PNs *in vivo* are largely unknown, preventing the exploration of its functional significance. Here, we report that the compartmentalized morphology of dendritic mitochondria characterizing CA1 PNs *in vivo* is present early in development, refined during postnatal maturation of CA1 PNs dendrites and requires activity-dependent and input-specific regulation of the Camkk2- AMPK kinase dyad. The relationship between activity and mitochondrial dynamics differs across the dendrites, as, in proximal dendrites, but not in apical tuft dendrites, high levels of activity-dependent Camkk2-mediated AMPK phosphorylates two direct effectors (Mff and Mtfr1l) that respectively promote mitochondrial fission and oppose fusion. Thus, our results identify a novel activity-dependent molecular pathway regulating the compartment-specific morphology of dendritic mitochondria of CA1 PNs *in vivo*.

## RESULTS

### CA1 hippocampal pyramidal neurons display sub-cellular, compartment specific dendritic mitochondria morphology *in vivo*

In order to visualize mitochondrial morphology in adult CA1 PNs *in vivo* in unfixed and fixed conditions, we used two complementary electroporation techniques: single cell electroporation (SCE; **Fig. 1B**) and *in utero* electroporation (IUE; **Fig. 1A**). Both techniques enable the labeling of mitochondria to visualize their morphology with enough sparsity for optical isolation of single CA1 PNs *in vivo*. The SCE approach has the advantage of enabling *in vivo* live 2-photon imaging of mitochondria in an anesthetized mouse, preventing any potential artifact of fixation which can affect mitochondria morphology ^26^. IUE also allows for sparse expression of mitochondrial markers *in vivo* and is coupled with fixation and imaging with confocal microscopy on brain slices. In both techniques, we co-electroporated a cytoplasmic fluorescent protein (tdTomato, mGreenLantern or mTagBFP2) and a mitochondrial matrix-targeted fluorescent protein (mt-YFP, mt-DsRed or mt-mTAGBFP2) allowing us to visualize single mitochondrial matrices in optically isolated dendrites in developing or adult CA1 PNs.

**Figure 1.**
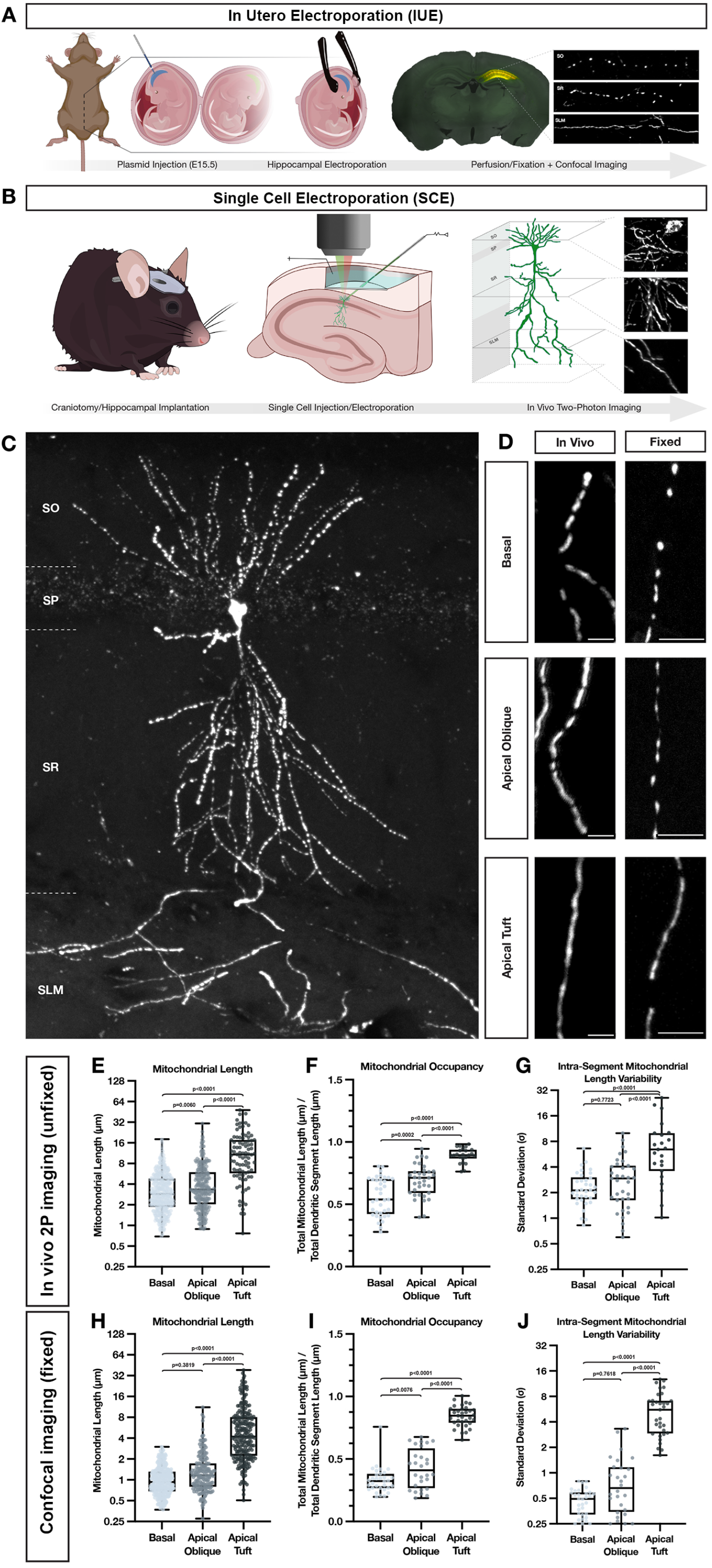
**Mitochondria display highly compartmentalized, layer-specific, morphology in dendrites of CA1 PNs *in vivo*.** (A) Schematic of *in utero* electroporation (IUE) procedure used to sparsely express fluorescent reporters, cDNAs or shRNAs in CA1 PNs of the hippocampus. (B) Schematic of single cell electroporation (SCE), used to express fluorescent reporters into single adult CA1 pyramidal neurons in the hippocampus. This method allows for rapid expression of plasmid DNA and *in vivo* two-photon visualization of individual neurons in an adult anesthetized mouse *in vivo*. (C) Representative image of a single CA1 PN expressing a mitochondrial matrix reporter (mt-YFP) via SCE following fixation and re-imaged using confocal microscopy on a single vibratome section. (D) High-magnification representative images of dendrites from the three hippocampal compartments—basal (SO), apical oblique (SR), and apical tuft (SLM)—expressing a mitochondrial matrix marker (mt-YFP or mt- DsRed) imaged either *in vivo* using two-photon microscopy (left) or post-fixation with confocal microscopy (right). Scale bar, 5 μm. (**E**-**G**) Quantification of individual mitochondrial length (**E**), mitochondrial segment occupancy (**F**), and intra-segment mitochondrial length variability (**G**) from basal, apical oblique, and apical tuft dendrites captured *in vivo*. (Basal: n = 422 mitochondria; n = 37 dendritic segments; mean length = 3.621 μm ± 0.124 (SEM); mean occupancy = 56.11% ± 2.49% (SEM); mean intra-segment variability = 2.420 μm ± 0.198 (SEM); Apical Oblique: n = 263 mitochondria; n = 35 dendritic segments; mean length = 4.719 μm ± 0.249 (SEM); mean occupancy = 68.82% ± 2.20% (SEM); mean intra-segment variability = 3.185 μm ± 0.370 (SEM); Apical Tuft: n = 90 mitochondria; n = 23 dendritic segments; mean length = 13.02 μm ± 1.041 (SEM); mean occupancy = 89.26% ± 1.24% (SEM); mean intra-segment variability = 8.018 μm ± 6.359 (SEM)). (**H**-**J**) Quantification of individual mitochondrial length (**H**), mitochondrial segment occupancy (**I**), and intra-segment mitochondrial length variability (**J**) from basal, apical oblique, and apical tuft dendrites following fixation. (Basal: n = 313 mitochondria; n = 32 dendritic segments; mean length = 1.065 μm ± 0.029 (SEM); mean occupancy = 33.01% ± 1.84% (SEM); mean intra-segment variability = 0.4736 μm ± 0.030 (SEM); Apical Oblique: n = 184 mitochondria; n = 28 dendritic segments; mean length = 1.569 μm ± 0.104 (SEM); mean occupancy = 42.38% ± 2.98% (SEM); mean intra-segment variability = 0.9169 μm ± 0.149 (SEM); Apical Tuft: n = 226 mitochondria; n = 33 dendritic segments; mean length = 6.562 μm ± 0.437 (SEM); mean occupancy = 84.19% ± 1.44% (SEM); mean intra-segment variability = 5.695 μm ± 0.552 (SEM)).

As previously reported ^11^, quantitative measurement of mitochondrial matrix size and dendritic occupancy following fixation of IUE CA1 PNs shows a remarkable degree of subcellular compartmentalization corresponding with spatially restricted afferents arriving onto distinct portions of the dendritic arbor (**Fig. 1C-D, H-J**). In basal dendrites (SO) of CA1 PNs located, mitochondria are short (1.065 μm ± 0.029), occupy a third of the dendritic segments (33.01% ± 1.84%), and are uniform in length (mean intra-segment variability: 0.4736 μm ± 0.030). In apical oblique dendrites located in SR, mitochondria are also short, though nearly 1.5 times longer than in SO (1.569 μm ± 0.104), occupy just under half of the dendritic processes (42.38% ± 2.98%), and are rather uniform (mean intra-segment variability: 0.9169 μm ± 0.149). In apical tuft dendrites located in SLM, mitochondria are significantly more elongated (6.562 μm ± 0.437) than in SO or SR, occupy a larger proportion of dendritic segments (84.19% ± 1.44%), and vary significantly more in size within individual dendritic segments (mean intra-segment variability, 5.695 μm ± 0.552).

To exclude the possibility that this compartment-specific morphology of dendritic mitochondria could be the result of fixation ^26^, we turned to SCE to image dendritic mitochondria of CA1 PNs in living mice. Two to three days following SCE with a cytoplasmic and mitochondrial matrix marker, mice were placed under isoflurane anesthesia while individual dendritic segments from all three compartments—basal, apical oblique, and apical tuft—were imaged using 2-photon microscopy. Despite different imaging conditions, quantitative measurements of size, occupancy, and intra-segment variability show conserved relationships between all three compartments with both approaches (**Fig. 1D, E-G**). These results demonstrate that the striking degree of compartmentalization of dendritic mitochondria characterizing CA1 PNs *in vivo* is not the result of fixation artifacts.

Because the above observations were made using mitochondrial matrix-targeted fluorescent reporters, we repeated the IUE experiments, this time expressing fluorescent proteins targeted to both the mitochondrial matrix (mt-mTAGBFP2 or mt-YFP) and the outer mitochondrial membrane (OMM; ActA-mCherry-HA). Quantification of both mitochondrial size and occupancy using both OMM and matrix markers confirmed the above observations. Interestingly, however, values obtained with the OMM marker were significantly longer in all three compartments than their respective matrix measurements, while still maintaining significantly compartmentalized differences (**Fig. S1A-C**).

This result indicates that individual mitochondria defined by OMM markers can contain fragmented matrix sub- volumes, which can be challenging to resolve using diffraction-limited light microscopy. Previous work has reported that in neuronal dendrites, the IMM can undergo repetitive constriction events, termed CoMIC, independently of OMM membrane dynamics ^27^. In order to determine the ultrastructural features of mitochondria in CA1 PNs dendrites *in vivo*, and to confirm if dendritic mitochondria in SLM and SR adopted distinct morphology with an independent imaging approach, we took advantage of a serial EM dataset previously published for connectomic analysis ^28^. In this dataset, mitochondria located in the apical dendrites of CA1 PNs located in SLM and SR were reconstructed in 3D (**Fig. S1D-H**) and our analysis confirms striking differences in mitochondria morphology between these two dendritic compartments. In SR, mitochondria adopt a unique morphology with thin constrictions (often <100nm in diameter; arrow in **Fig. S1E**) and bulging of the matrix (arrowheads in **Fig. S1E**). This unique ultrastructural feature is seen much less frequently in mitochondria found in SLM which presents a more uniformly tubular morphology (see multiple examples in **Fig. S1H**). This morphological difference results in a significant reduction in the volume of dendrite occupied by mitochondria in SR compared to SLM (**Fig. S1I**).

Overall, our light and electron microscopy approaches converge to show that mitochondrial morphology differs significantly in specific dendritic compartments of CA1 PNs *in vivo* with highly elongated and tubular morphology in SLM and progressively more fragmented matrix compartments in SR and SO.

### Compartment-specific dendritic mitochondria morphology is present early in the development of CA1 PNs *in vivo* but is absent *in vitro*

To determine the developmental timeframe for the appearance of compartmentalized mitochondria morphology described above in adult CA1 PNs *in vivo,* we collected *in utero* electroporated brains at postnatal days seven, ten, fourteen and twenty-one (P7, P10, P14 and P21). At all stages of development, we observed that mitochondria are significantly shorter in the basal dendrites located in SO than in the apical tuft dendrites located in SLM (**Fig. S2A**). Interestingly, while this compartment-specific mitochondrial morphology is clearly present at each developmental timepoint, mitochondria increase in both size (**Fig. S2B**) and dendritic occupancy (**Fig. S2C**) in both domains throughout maturation and reach adult-like values by P21.

To determine whether compartmentalized mitochondrial morphology characterizing CA1 PNs *in vivo* is regulated by cues intrinsic to the dendrites or cues extrinsic to the cell, we performed IUE to label mitochondria of CA1 PNs at E15.5 as above but at E18.5, the hippocampi of electroporated embryos were collected, enzymatically dissociated, and maintained in 2-dimensional (2D) cultures *in vitro* (**Fig. S3A-B**). Following culture for ten, fourteen or eighteen days in vitro (DIV10, 14, 18), we observed either no significant difference in mitochondrial length or occupancy (10, 18DIV), or opposite results to those found *in vivo*: mitochondria were slightly but significantly longer with higher occupancy values in the proximal dendrites compared to distal dendrites (14DIV) (**Fig. S3C-D**). These results suggest that factors extrinsic to the cell, including local activity patterns, might drive the formation of the compartment specific mitochondrial morphology observed *in vivo*.

### Neuronal activity and domain-specific synaptic inputs regulate the formation of compartmentalized mitochondrial morphology of CA1 PNs *in vivo*

Previous results obtained *in vitro* suggest that synaptic activity induces local Drp1-dependent mitochondrial fission events in the context of synaptic plasticity induction ^12^. To test if the compartmentalized morphology of dendritic mitochondria observed in CA1 PNs *in vivo* is regulated by neuronal activity, we first performed IUE to express the inward rectifying potassium channel 2.1 (Kir2.1) which hyperpolarizes neurons to ∼-90mV and drastically reduces their excitability and ability to fire action potentials ^29, 30^. Strikingly, Kir2.1 overexpression in CA1 PNs *in vivo* through development strongly reduced the compartment-specific differences in mitochondria morphology. Dendritic mitochondria in Kir2.1 expressing CA1 PNs became significantly longer and occupied a higher percentage of the dendrites in SO (basal dendrites, **Fig. 2A-C**), SR (apical obliques, **Fig. 2D-F**), and SLM (apical tufts, **Fig. 2G-I**) compared to control. This result demonstrates that neuronal activity is required for the proper maturation of compartmentalized morphology of dendritic mitochondria in CA1 PNs *in vivo*.

**Figure 2.**
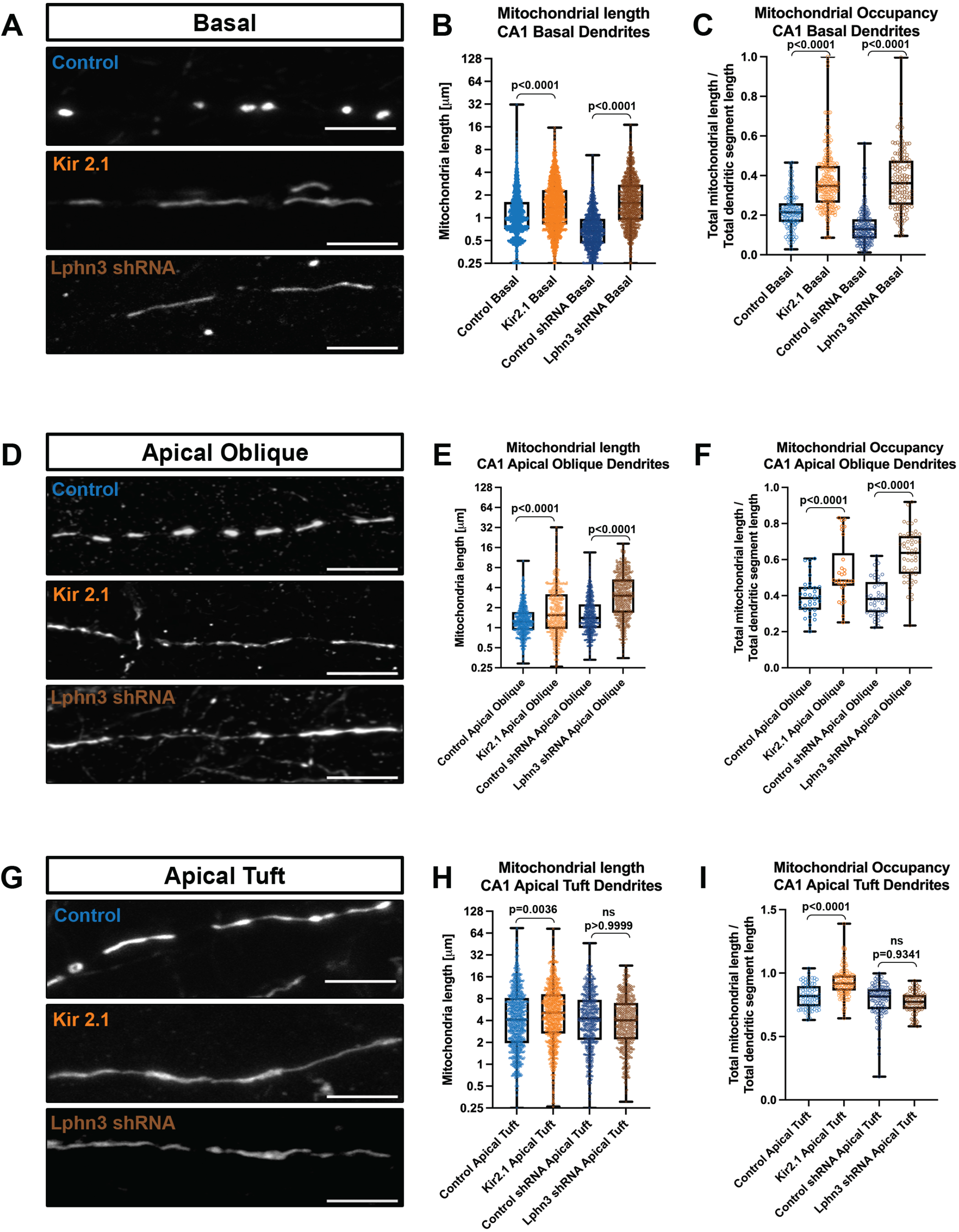
Neuronal activity regulates mitochondrial size in a compartment specific manner in CA1 pyramidal neurons *in vivo*. (**A-I**) High magnification representative images of mitochondrial morphology within isolated secondary or tertiary hippocampal CA1 (**A**) basal, (**D**) apical oblique, and (**G**) apical tuft dendrites in which a mitochondrial matrix-targeted fluorescent protein (mt-YFP) was IUE along with either a control plasmid (pCAG tdTomato) or pCAG Kir2.1-T2A-tdTomato), or either control shRNA or Lphn3 shRNA plasmid. Quantification of mitochondrial length and occupancy in the (**B-C**) basal, (**E-F**) apical oblique, and (**H-I**) apical tuft dendritic compartments following Kir2.1 over-expression or shRNA-mediated knockdown of Lphn3. Quantification of mitochondrial length (**B**, **E,** and **H**) and mitochondrial occupancy (**C**, **F** and **I**) in basal dendrites (**B-C**), apical oblique (**E-F**) and apical tuft (**H-I**) demonstrates that both Kir2.1 over-expression and Lphn3 knockdown significantly increases mitochondrial length and occupancy in basal (**B-C**) and apical oblique dendrites (**E-F**). Control_tdTomato-basal_ = 127 segments, 2154 mitochondria, mean length = 1.42µm ± 0.03µm (SEM), mean occupancy = 21.8% ± 0.9% (SEM); Control_tdTomato-apical oblique_ = 34 segments, 748 mitochondria, mean length = 1.49µm ± 0.04µm, mean occupancy = 39.4% ± 1.0%; Control_tdTomato-apical tuft_ = 75 segments, 1036 mitochondria, mean length = 6.48µm ± 0.23µm, mean occupancy = 81.9% ± 1.1%; Kir2.1_tdTomato-basal_ = 206 segments, 2278 mitochondria, mean length = 1.87µm ± 0.03µm, mean occupancy = 37.8% ± 1.0%; Kir2.1_tdTomato-apical oblique_ = 30 segments, 435 mitochondria, mean length = 2.54µm ± 0.14µm, mean occupancy = 53.5% ± 2.9%; Kir2.1_tdTomato-apical tuft_ = 128 segments, 965 mitochondria, mean length = 7.19µm ± 0.22µm, mean occupancy = 92.7% ± 1.0%; Control_shRNA-basal_ = 169 segments, 1344 mitochondria, mean length = 0.81µm ± 0.01µm, mean occupancy = 13.8% ± 0.6%; Control_shRNA-apical oblique_ = 45 segments, 633 mitochondria, mean length = 1.84µm ± 0.05µm, mean occupancy = 39% ± 1.6%; Control_shRNA-apical tuft_ = 106 segments, 617 mitochondria, mean length = 5.81µm ± 0.20µm, mean occupancy = 78.6% ± 1.2%; Lphn3_shRNA-basal_ = 129 segments, 1169 mitochondria, mean length = 2.11µm ± 0.04µm, mean occupancy = 37.4% ± 1.3%; Lphn3_shRNA-apical oblique_ = 53 segments, 573 mitochondria, mean length = 3.92µm ± 0.12µm, mean occupancy = 62.5% ± 1.9%; Lphn3_shRNA-apical tuft_ = 84 segments, 561 mitochondria, mean length = 5.1µm ± 0.16µm, mean occupancy = 75.2% ± 0.9%. p values are indicated in the figure a following Kruskal-Wallis tests. Data are shown as individual points on box plots with 25^th^, 50^th^ and 75^th^ percentiles indicated with whiskers indicating min and max values. Scale bar, 5 μm.

We next perturbed the synaptic inputs received by CA1 PNs in compartment-specific manner by downregulating Latrophilin 3 (Lphn3, **Fig. S4**), a postsynaptic adhesion molecule recently shown to be required for the ability of CA3 axons to form ∼50% of synapses onto dendrites of CA1 PNs specifically in the SO and SR compartments, but not for synapses made by EC axons in SLM ^31^. This synapse-specific manipulation results in a reduction by ∼50% of miniature excitatory potential synaptic currents (mEPSC) in CA1 PNs ^31^. We confirmed that shRNA-mediated knockdown of Lhpn3 (**Fig. S4A-B**) by IUE of CA1 PNs leads to ∼40% reduction in spine density in SO but not in SLM (**Fig. S4C-D**).

Interestingly, this reduction of presynaptic input mediated by Lphn3 knockdown in CA1 PNs led to a striking and significant increase in mitochondria size and occupancy in dendritic compartments receiving CA3 inputs (SO and SR; **Fig. 2A-F**), while mitochondria morphology in the apical tuft (SLM) of the same CA1 PNs were unaffected (**Fig. 2G-I**). These results show that (1) local synaptic activity is a major regulator of mitochondria morphology in dendrites of CA1 PNs and (2) suggests that an activity-dependent signaling mechanism promotes the small mitochondria morphology in SO and SR *in vivo,* potentially by promoting mitochondria fission and/or inhibiting mitochondria fusion in basal and apical dendrites but not in the apical tuft.

### Compartmentalized mitochondrial morphology in CA1 PNs requires Camkk2 and AMPK *in vivo*

We and others previously showed that the kinase dyad Camkk2 and AMPK are regulated by neuronal activity since both neuronal depolarization, activation of VGCCs by depolarization, or activation of NMDA receptors can activate AMPK in a Camkk2-dependent manner in cortical and hippocampal neurons ^24, 32^. Furthermore, we also demonstrated that Camkk2-dependent AMPK over-activation mediates excessive mitochondrial fission through direct phosphorylation of Mff in apical dendrites of CA1 PNs ^11, 23^. Therefore, we hypothesized that synaptic activity could regulate mitochondrial compartmentalization in a Camkk2-AMPK-dependent manner whereby low Camkk2-AMPK activity in apical tufts of CA1 PNs leads to fusion-dominant mitochondrial elongation in SLM and higher activity in basal and apical oblique dendrites leads to fission-dominant smaller mitochondrial morphology in SO and SR.

To test this in CA1 PNs *in vivo*, we performed CA1-targeted IUE in a constitutive *Camkk2* knockout mouse, as well as in a conditional *AMPK*α*1*^F/F^α*2*^F/F^ mouse (AMPK cDKO) and measured mitochondrial length and occupancy using a mitochondrial matrix-targeted fluorescent protein (mt-YFP, mt-DsRed) (**Fig. 3**). By electroporating Cre recombinase with a Cre-dependent reporter (FLEX-tdTomato, FLEX-mGreenLantern) in the AMPK cDKO, we could delete AMPK only from a subset of CA1 PNs. Using these approaches, we found that AMPK-null CA1 PNs display significantly increased mitochondria length and occupancy in basal (SO, **Fig. 3A-C**) and apical oblique (SR, **Fig. 3D-F**), but not in the apical tuft (SLM, **Fig. 3G-I**) dendritic compartments. Similarly, Camkk2-null CA1 PNs show a remarkably similar increase in mitochondrial size and occupancy in both SO and SR dendritic compartments (**Fig. 3A-F**). One interesting difference is that Camkk2-null CA1 PNs display a small but significant increase in mitochondria size and occupancy in the apical tuft compartment (SLM) not observed in the AMPK-null CA1 PNs. These results overall demonstrate that Camkk2 and AMPK are both required for the compartmentalized morphology of dendritic mitochondria observed *in vivo* and are specifically required for the small mitochondria size and dendritic occupancy in SO and SR dendritic compartments of CA1 PNs *in vivo*.

**Figure 3.**
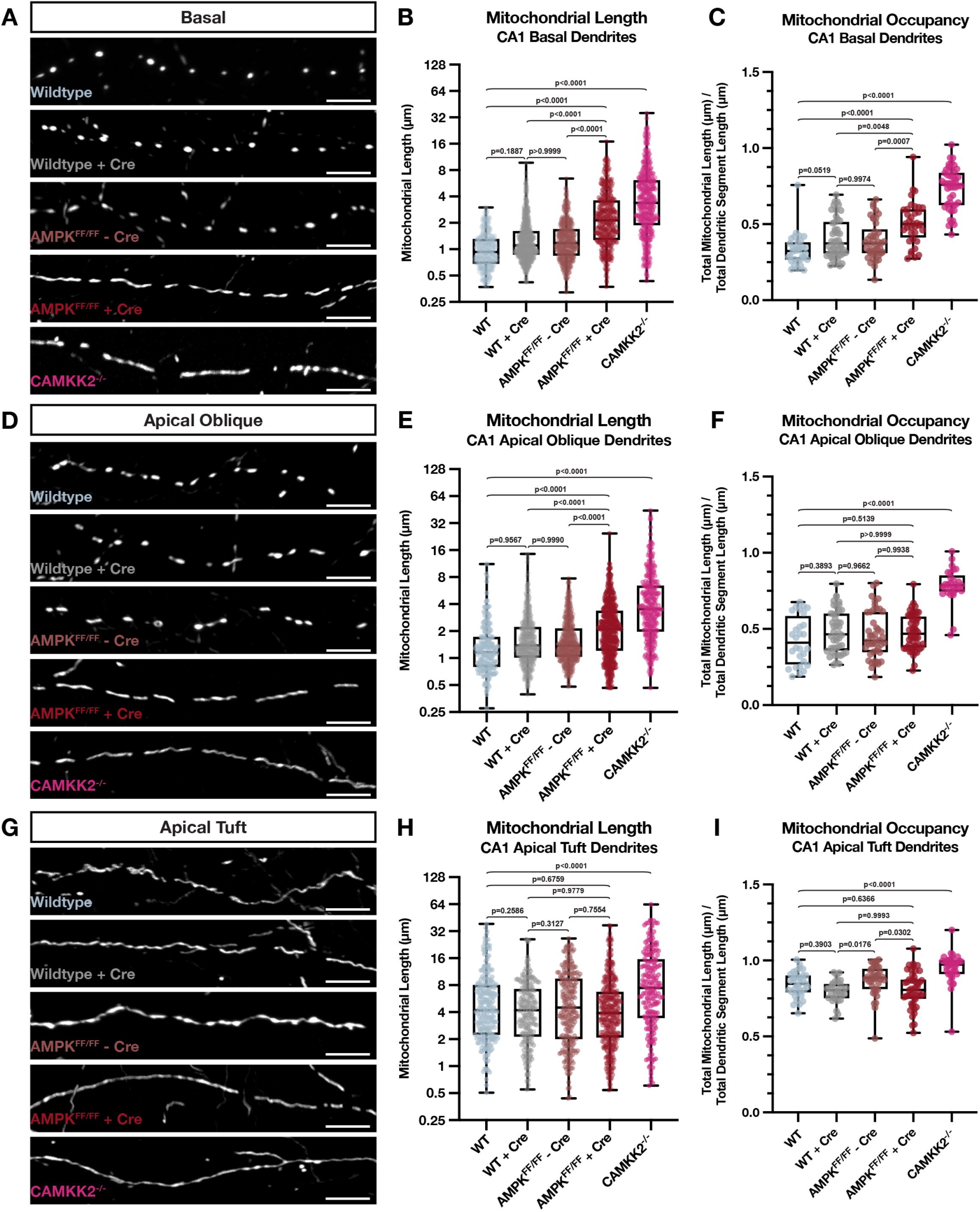
Camkk2 and AMPK are required for the compartment-specific mitochondrial morphology in CA1 PNs *in vivo*. (**A-I**) High magnification representative images of mitochondrial morphology within isolated secondary or tertiary hippocampal CA1 (**A**) basal, (**D**) apical oblique, and (**G**) apical tuft dendrites in which a mitochondrial matrix-targeted fluorescent protein (mt-YFP or mt-DsRed) and cell fill (tdTomato or mGreenLantern—not pictured) were co-expressed by IUE with or without Cre recombinase (+/-Cre). AMPKα1^F/F^α2^F/F^ double conditional mice were *in utero* electroporated with the same mitochondrial markers/cell fills and either no Cre (AMPK^FF/FF^ - Cre) or Cre (AMPK^FF/FF^ + Cre). Camkk2^-/-^ constitutive knock-out mice were electroporated with the same mitochondrial markers/cell fills as above (Camkk2^-/-^). Quantification of mitochondrial length and occupancy in the (**B-C**) basal, (**E-F**) apical oblique, and (**H-I**) apical tuft dendritic compartments reveals a significant increase in mitochondrial length (**B**) and occupancy (**C**) in basal dendrites when knocking out AMPK or CAMKK2 when compared to their controls, with Camkk2 KO having a much bigger effect (WT: n = 32 dendritic segments, 313 individual mitochondria, mean length = 1.065 μm ± 0.029 (SEM), mean occupancy = 33.01% ± 1.84%; WT + Cre: n = 46 dendritic segments, 627 individual mitochondria, mean length = 1.437 μm ± 0.040 (SEM), mean occupancy = 40.95% ± 1.94%; AMPK^FF/FF^ - Cre: n = 42 dendritic segments, 463 individual mitochondria, mean length = 1.418 μm ± 0.039 (SEM), mean occupancy = 39.24% ± 1.89%; AMPK^FF/FF^ + Cre: n = 36 dendritic segments, 381 individual mitochondria, mean length = 2.832 μm ± 0.116 (SEM), mean occupancy = 50.72% ± 2.36%, length increase = 97.08%, occupancy increase = 23.86%; Camkk2^-/-^: n = 46 dendritic segments, 449 individual mitochondria, mean length = 4.975 μm ± 0.233 (SEM), mean occupancy = 73.36% ± 2.03%, length increase = 367%, occupancy increase = 122%). A significant increase in mitochondrial length (**E**) and occupancy (**F**) is also seen in apical oblique dendrites when knocking out Camkk2, but only an increase in length is seen when knocking out AMPK. (WT: n = 28 dendritic segments, 184 individual mitochondria, mean length = 1.569 μm ± 0.104 (SEM), mean occupancy = 42.38% ± 2.98%; WT + Cre: n = 44 dendritic segments, 525 individual mitochondria, mean length = 1.791 μm ± 0.052 (SEM), mean occupancy = 48.50% ± 2.12%; AMPK^FF/FF^ - Cre: n = 45 dendritic segments, 440 individual mitochondria, mean length = 1.885 μm ± 0.075 (SEM), mean occupancy = 45.82% ± 2.29%; AMPK^FF/FF^ + Cre: n = 52 dendritic segments, 499 individual mitochondria, mean length = 2.645 μm ± 0.098 (SEM), mean occupancy = 47.79% ± 1.62%, length increase = 47.68%; Camkk2^-/-^: n = 29 dendritic segments, 288 individual mitochondria, mean length = 5.366 μm ± 0.346 (SEM), mean occupancy = 78.64% ± 2.07%, length increase = 242%, occupancy increase = 85.56%). Note that in Camkk2-null CA1 PNs we observe a significant increase in mitochondrial length (**H**) and occupancy (**I**) in the apical tuft compared to WT controls (WT: n = 33 dendritic segments, 226 individual mitochondria, mean length = 6.562 μm ± 0.437 (SEM), mean occupancy = 84.19% ± 1.44%; WT + Cre: n = 29 dendritic segments, 189 individual mitochondria, mean length = 5.206 μm ± 0.052 (SEM), mean occupancy = 79.40% ± 1.38%; AMPK^FF/FF^ - Cre: n = 38 dendritic segments, 229 individual mitochondria, mean length = 6.498 μm ± 0.368 (SEM), mean occupancy = 87.00% ± 1.67%; AMPK + Cre: n = 47 dendritic segments, 370 individual mitochondria, mean length = 5.723 μm ± 0.300 (SEM), mean occupancy = 80.65% ± 1.71%; Camkk2^-/-^: n = 35 dendritic segments, 199 individual mitochondria, mean length = 11.14 μm ± 0.758 (SEM), mean occupancy = 95.03% ± 1.73%, length increase = 69.77%, occupancy increase = 12.88%). Scale bar, 5 μm.

### Compartmentalized mitochondrial morphology in CA1 PNs requires Mtfr1l-dependent inhibition of Opa1

Previous results identified two direct effectors phosphorylated by AMPK mediating mitochondrial fission: the Drp1-receptor Mff ^23^ and the anti-fusion, Opa1-inhibiting protein Mitochondrial Fission Regulator 1-Like (Mtfr1l)^33^. We performed CA1-targeted IUE with cytoplasmic fluorescent proteins (tdTomato, mGreenLantern) and mitochondrial matrix-targeted fluorescent proteins (mt-YFP, mt-DsRed) in combination with either a control, non-targeting shRNA (shNT) or an shRNA targeting *Mtfr1l* (**Fig. 4**). The shRNA selected to knockdown Mtfr1l is validated in **Fig. S5A-B**. We found that knocking down Mtfr1l in CA1 PNs significantly elongates mitochondria and increases mitochondrial occupancy in the basal (SO; **Fig. 4A-C**) and apical oblique (SR; **Fig. 4D-F**) dendritic compartments, but not in apical tuft (SLM) dendrites (**Fig. 4H**). To confirm the specificity of our *Mtfr1l*- targeting shRNA, we co-expressed it together with a shRNA-resistant human cDNA expressing hMTFR1L. We found, as expected, that expression of hMTFR1L is sufficient to rescue mitochondrial length (**Fig. 4B,E**) and occupancy (**Fig. 4C,F**) back to levels observed in SO and SR dendritic compartments of control shRNA expressing CA1 PNs. Recent work demonstrated that Mtfr1l exerts its effects on mitochondria size by inhibiting the pro-fusion protein Opa1, and that functionally, Mtfr1l represents a novel class of ‘anti-fusion’ mitochondrial proteins because of its ability to oppose fusion and thereby enable Mff-dependent fission ^33^. Based on these previously published results, we hypothesized that the mitochondrial elongation mediated by knocking down Mtfr1l should be rescued by simultaneously knocking down Opa1 in CA1 PNs. Therefore, we performed IUEs with shRNAs targeting both *Mtfr1l* and *Opa1* (see validation in **Fig. S5E-F**). Knocking down Opa1 completely rescues the effect of knocking down Mtfr1l and re-establishes short mitochondria and low occupancy in both basal and apical oblique dendrites to levels observed in control CA1 PNs (**Fig. 4A-F**) No significant effects were observed in dual knockdown of Mtfr1l and Opa1 in the apical tuft of the same neurons compared to knockdown of Mtfr1l alone or control shRNA expressing neurons (**Fig. 4H,I**). Overall, these results demonstrate that Mtfr1l is required for the compartmentalized morphology of mitochondria observed in the basal and apical oblique dendritic domains of CA1 PNs *in vivo* through inhibition of Opa1.

**Figure 4.**
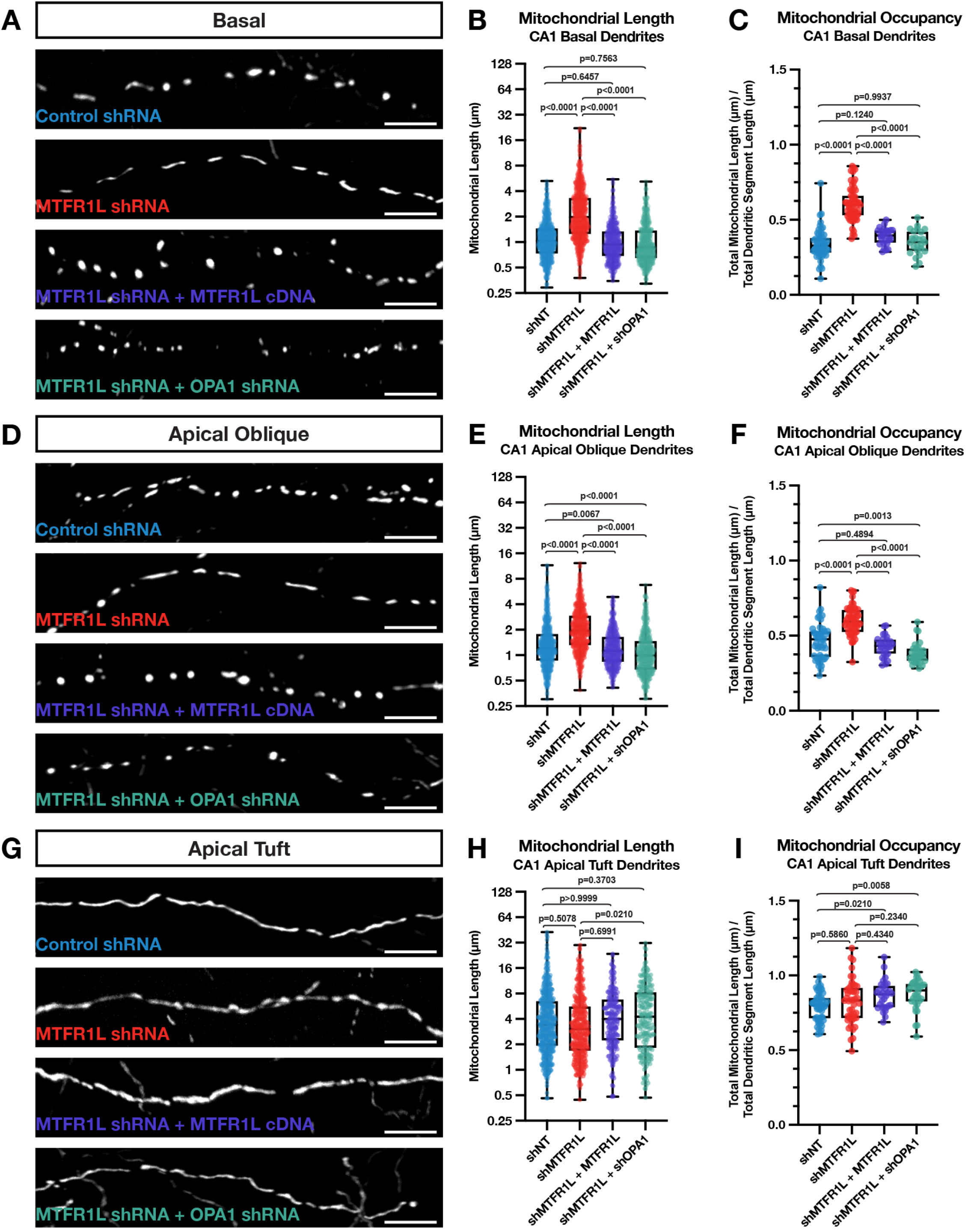
MTFR1L restricts hippocampal CA1 PN basal and apical oblique dendritic morphology through inhibition of Opa1. High magnification representative images of mitochondrial morphology within isolated secondary or tertiary hippocampal CA1 **A**) basal, **D**) apical oblique, and **G**) apical tuft dendrites in which a mitochondrial matrix-targeted fluorescent protein (mt-YFP or mt-DsRed) and cell fill (tdTomato or mGreenLantern—not pictured) was *in utero* electroporated along with either a control shRNA (shNT), *Mtfr1l* shRNA (shMtfr1l), *Mtfr1l* shRNA with full-length hMTFR1L cDNA (shMtfr1l + hMTFR1L), or *Mtfr1l* shRNA and *Opa1* shRNA (shMtfr1l + shOpa1). Quantification of mitochondrial length and occupancy in the (**B-C**) basal, (**E-F**) apical oblique, and (**H-I**) apical tuft dendritic compartments reveals a significant increase in mitochondrial length (**B**) and occupancy (**C**) in basal dendrites when knocking down Mtfr1l, which is rescued when re-expressing full-length hMTFR1L or knocking down Opa1 (shNT: n = 51 dendritic segments, 670 individual mitochondria, mean length = 1.225 μm ± 0.028 (SEM), mean occupancy = 33.95% ± 1.46%; shMTFR1L: n = 60 dendritic segments, 773 individual mitochondria, mean length = 2.721 μm ± 0.085 (SEM), mean occupancy = 60.36% ± 1.46%, length increase = 122%, occupancy increase = 77.79%; shMtfr1l + hMTFR1L: n = 28 dendritic segments, 511 individual mitochondria, mean length = 1.113 μm ± 0.029 (SEM), mean occupancy = 39.10% ± 1.02%; shMtfr1l + shOpa1: n = 25 dendritic segments, 515 individual mitochondria, mean length = 1.126 μm ± 0.032 (SEM), mean occupancy = 35.08% ± 1.61%). A significant increase in mitochondrial length (**E**) and occupancy (**F**) is also seen in apical dendrites when knocking down Mtfr1l, which is similarly rescued when either expressing full-length hMTFR1L or additionally knocking down Opa1 (shNT: n = 48 dendritic segments, 669 individual mitochondria, mean length = 1.584 μm ± 0.059 (SEM), mean occupancy = 46.34% ± 1.72%; shMtfr1l: n = 54 dendritic segments, 683 individual mitochondria, mean length = 2.390 μm ± 0.061 (SEM), mean occupancy = 59.86% ± 1.29%, length increase = 50.88%, occupancy increase = 29.18%; shMtfr1l + hMTFR1L: n = 29 dendritic segments, 490 individual mitochondria, mean length = 1.352 μm ± 0.034 (SEM), mean occupancy = 42.92% ± 1.25%; shMtfr1l + shOpa1: n = 32 dendritic segments, 514 individual mitochondria, mean length = 1.248 μm ± 0.038 (SEM), mean occupancy = 38.28% ± 1.23%). There’s no significant effect of knocking down Mtfr1l in the apical tuft dendrites in either mitochondrial length (**H**) or occupancy **(I**) compared to control neurons (shNT: n = 53 dendritic segments, 472 individual mitochondria, mean length = 5.177 μm ± 0.250 (SEM), mean occupancy = 79.14% ± 1.29%; shMtfr1l: n = 46 dendritic segments, 388 individual mitochondria, mean length = 4.665 μm ± 0.236 (SEM), mean occupancy = 82.35% ± 2.17%; shMtfr1l + hMTFR1L: n = 30 dendritic segments, 219 individual mitochondria, mean length = 5.186 μm ± 0.280 (SEM), mean occupancy = 86.65% ± 1.75%; shMtfr1l + shOpa1: n = 34 dendritic segments, 215 individual mitochondria, mean length = 5.872 μm ± 0.352 (SEM), mean occupancy = 87.36% ± 1.72%). Scale bar, 5 μm

### Compartmentalized mitochondrial morphology in CA1 PNs requires Mff

AMPK has also been shown to regulate mitochondrial morphology through direct phosphorylation of the OMM localized Drp1 receptor Mff ^19, 23^. To test if Mff is required for the compartmentalized morphology of mitochondria in dendrites of CA1 PNs, we repeated the same IUE experiments using an shRNA targeting *Mff* (shMff) previously characterized ^7^, and validated the shRNA efficiently knocks down all isoforms of Mff (**Fig. S5C-D**). Upon knocking down Mff in CA1 PNs using IUE, we observe a significant increase in mitochondrial length and occupancy in all three dendritic compartments (**Fig. 5A-I**). Since knockdown of either Mtfr1l or Mff leads to similar phenotypes on mitochondria morphology, especially in basal and apical oblique dendritic domains of CA1 PNs, we tested if these two proteins have redundant or only partially overlapping functions in regulating dendritic mitochondrial morphology in these neurons. Interestingly, we found that knocking down both Mtfr1l and Mff does not lead to additive effects compared to knocking down Mff or Mtfr1l separately (**Fig. 5A-F**), strongly arguing that they play redundant functions or that they regulate their mutual abundance at OMM. In fact, we discovered that knockdown of either Mff or Mtfr1l significantly decreased protein abundance of the other; however, knockdown of Mff had a stronger impact on Mtfr1l expression (**Fig. S5G-H**). These results explain the observations that Mff knockdown has a greater effect on mitochondrial compartmentalization *in vivo*, and that no synergistic effect was seen upon simultaneous knockdown of both Mff and Mtfr1l.

**Figure 5.**
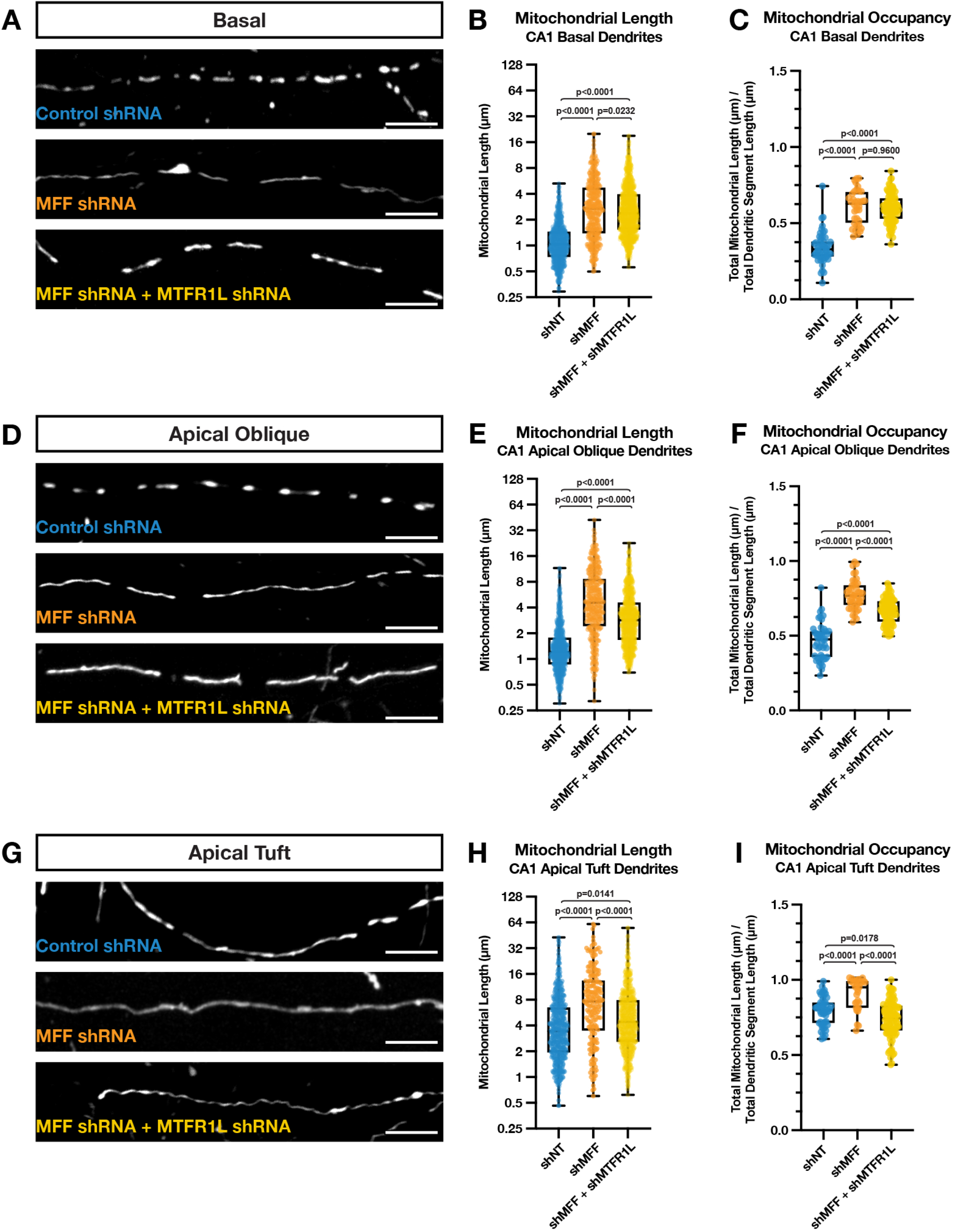
Mff restricts hippocampal CA1 PN basal, apical oblique, and apical tuft dendritic morphology. High magnification representative images of mitochondrial morphology within isolated secondary or tertiary hippocampal CA1 **A**) basal, **D**) apical oblique, and **G**) apical tuft dendrites in which a mitochondrial matrix-targeted fluorescent protein (mt-YFP or mt-DsRed) and cell fill (tdTomato or mGreenLantern—not pictured) was *in utero* electroporated along with either a control shRNA (shNT), *Mff* shRNA (shMFF), or *Mff* and *Mtfr1l* shRNA (shMff + shMtfr1l). Quantification of mitochondrial length and occupancy in the (**B-C**) basal, (**E-F**) apical oblique, and (**H-I**) apical tuft dendritic compartments reveals a significant increase in mitochondrial length (**B**) and occupancy (**C**) in basal dendrites when knocking down Mff, which does not show an additive effect when also knocking down Mtfr1l (shNT: n = 51 dendritic segments, 670 individual mitochondria, mean length = 1.225 μm ± 0.028 (SEM), mean occupancy = 33.95% ± 1.46%; shMff: n = 36 dendritic segments, 380 individual mitochondria, mean length = 3.530 μm ± 0.152 (SEM), mean occupancy = 60.83% ± 1.86%, length increase = 188%, occupancy increase = 79.18%; shMff + shMtfr1l: n = 92 dendritic segments, 1004 individual mitochondria, mean length = 3.180 μm ± 0.078 (SEM), mean occupancy = 59.94% ± 0.99%, length increase = 160%, occupancy increase = 76.55%). A significant increase in mitochondrial length (**E**) and occupancy (**F**) is also seen in apical dendrites when knocking down Mff, but to a significantly less degree when knocking down both Mff and Mtfr1l (shNT: n = 48 dendritic segments, 669 individual mitochondria, mean length = 1.584 μm ± 0.059 (SEM), mean occupancy = 46.34% ± 1.72%; shMff: n = 44 dendritic segments, 371 individual mitochondria, mean length = 6.445 μm ± 0.300 (SEM), mean occupancy = 77.34% ± 1.46%, length increase = 307%, occupancy increase = 66.90%; shMff + shMtfr1l: n = 85 dendritic segments, 829 individual mitochondria, mean length = 3.728 μm ± 0.106 (SEM), mean occupancy = 66.63% ± 0.93%, length increase = 135%, occupancy increase = 43.79%). While there is a significant increase mitochondrial length (**H**) and occupancy (**I**) in the apical tuft when knocking down Mff alone, this increase largely goes away when knocking down both Mff and Mtfr1l (shNT: n = 53 dendritic segments, 472 individual mitochondria, mean length = 5.177 μm ± 0.250 (SEM), mean occupancy = 79.14% ± 1.29%; shMff: n = 31 dendritic segments, 199 individual mitochondria, mean length = 10.78 μm ± 0.757 (SEM), mean occupancy = 90.44% ± 1.86%, length increase = 109%, occupancy increase = 14.28%; shMff + shMtfr1l: n = 95 dendritic segments, 708 individual mitochondria, mean length = 6.326 μm ± 0.236 (SEM), mean occupancy = 73.79% ± 1.27%, length increase = 22.19%). Scale bar, 5 μm.

### Camkk2 regulates morphological compartmentalization of dendritic mitochondria in CA1 PNs through AMPK-dependent Mtfr1l phosphorylation

We next tested whether the kinase dyad Camkk2-AMPK regulates the compartment-specific morphology of mitochondria in CA1 PNs through phosphorylation of Mtfr1l. We used two experimental approaches to test this. First, we performed biochemical analysis in mouse hippocampal neurons *in vitro* to test if Mtfr1l phosphorylation by AMPK is regulated by neuronal activity, and, whether activity-dependent regulation of Mtfr1l phosphorylation is Camkk2-dependent. To this end, we treated hippocampal neurons maintained in dissociated cultures for 3 weeks to a high potassium chloride (KCl) concentration (switch from 5 to 40mM) for a short time period (15min) to induce strong membrane depolarization and effectively open VGCCs leading to increased intracellular Ca^2+^ in hippocampal neurons ^34^. As shown previously ^24, 32^, western blot analysis revealed that neuronal depolarization induces increased AMPK phosphorylation on T172 in a Camkk2- dependent manner since it was blocked by a Camkk2-specific inhibitor STO609 ^24, 32^ (**Fig. 6A**). In the same lysates, KCl-mediated neuronal depolarization induced a significant increase in Mtfr1l phosphorylation on S103, one of the two serine residues phosphorylated by AMPK ^33^ that was also blocked by STO609 (**Fig. 6A**).

**Figure 6.**
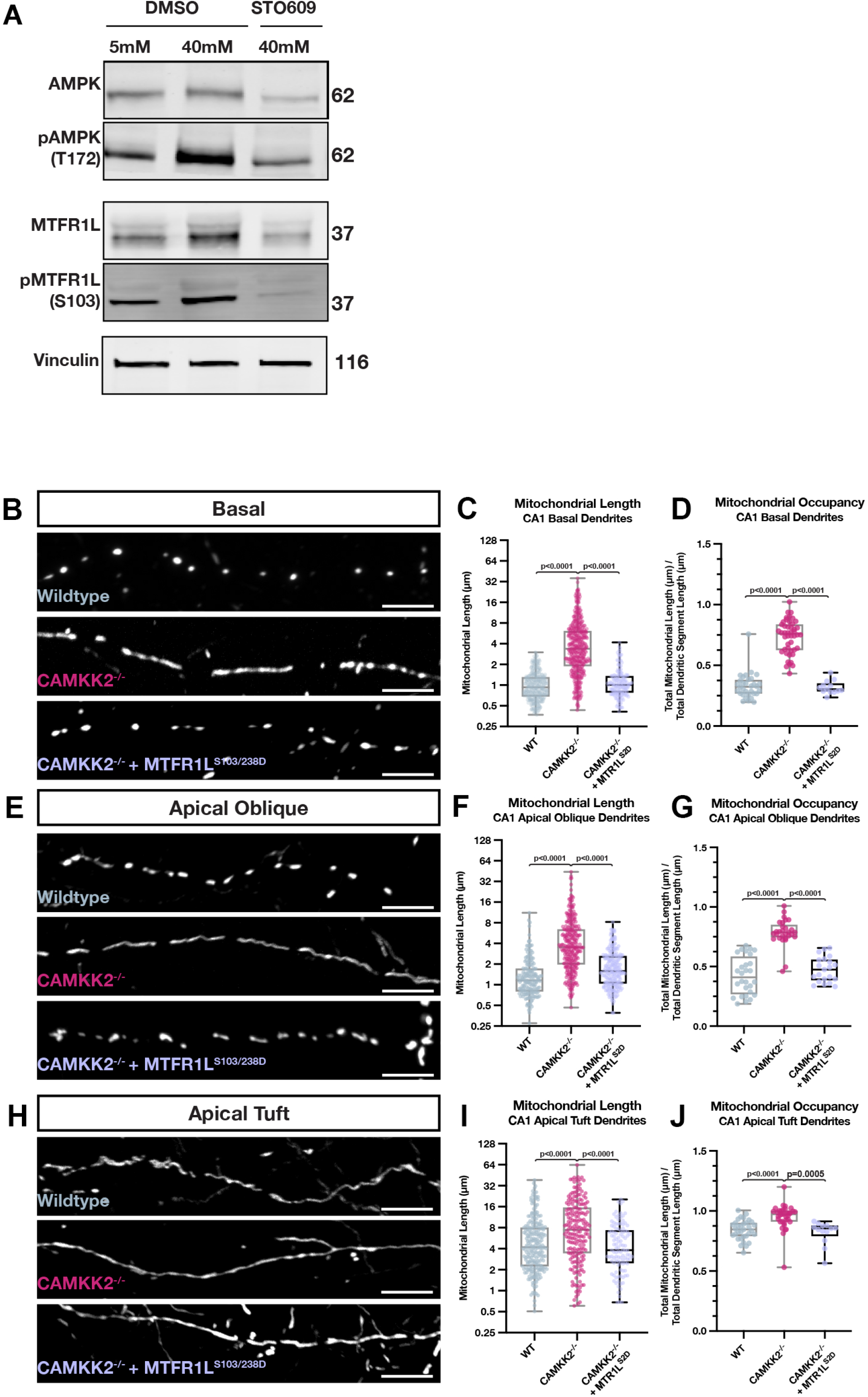
Activity-dependent and Camkk2-dependent phosphorylation of MTFR1L by AMPK mediates compartmentalized mitochondria morphology in dendrites of CA1 PNs *in vivo*. (A) Western blots (A) of whole cell lysates from mouse hippocampal neurons maintained in culture for 18- 21DIV and treated for 5 min with physiological (5mM) extracelllular potassium chloride (KCl) (first column) or high KCl (40mM) inducing membrane depolarization (columns 2 & 3) in the presence (column 3) or absence (columns 1&2) of the Camkk2 inhibitor STO609. These results demonstrate that phosphorylation of AMPKα catalytic subunit on T172 is increased by neuronal depolarization which is blocked by STO609. In turn, AMPK phosphorylation of its substrate MTFR1L on S103 ^33^ is increased by depolarization which is Camkk2- dependent since it is blocked by STO609. (B-J) Rescue experiments showing that phosphomimetic form of Mtfr1l (Mtfr1l^S2D^) mimicking phosphorylation by AMPK ^33^ is sufficient to rescue compartmentalized mitochondria morphology in basal (B-D), apical oblique (E-G) and apical tufts (H-J) dendrites of CA1 PNs *in vivo*. CA1 PNs from wild-type (WT) or *Camkk2*^-/-^ constitutive knockout mice were IUE with the same mitochondrial markers/cell fills as in Figure 3 (WT and Camkk2^-/-^) and a plasmid cDNA expressing phosphomimetic mutant on the two serine residues phosphorylated by AMPK (S103D and S238D) of Mtfr1l (Mtfr1l^S2D^) ^33^. Data and quantifications from WT and Camkk2^-/-^ are the same as in Figure 3. Basal Dendrites (WT: n = 32 dendritic segments, 313 individual mitochondria, mean length = 1.065 μm ± 0.029 (SEM), mean occupancy = 33.01% ± 1.84%; Camkk2^-/-^: n = 46 dendritic segments, 449 individual mitochondria, mean length = 4.975 μm ± 0.233 (SEM), mean occupancy = 73.36% ± 2.03%, length increase = 367%, occupancy increase = 122%; Camkk2^-/-^ + Mtfr1l^S2D^: n = 13 dendritic segments, 171 individual mitochondria, mean length = 1.187 μm ± 0.051 (SEM), mean occupancy = 31.89% ± 1.49%) Apical Oblique Dendrites (WT: n = 28 dendritic segments, 184 individual mitochondria, mean length = 1.569 μm ± 0.104 (SEM), mean occupancy = 42.38% ± 2.98%; Camkk2^-/-^: n = 29 dendritic segments, 288 individual mitochondria, mean length = 5.366 μm ± 0.346 (SEM), mean occupancy = 78.64% ± 2.07%, length increase = 242%, occupancy increase = 85.56%; Camkk2^-/-^ + Mtfr1l^S2D^: n = 21 dendritic segments, 223 individual mitochondria, mean length = 2.030 μm ± 0.092 (SEM), mean occupancy = 47.45% ± 2.28%) Apical Tuft Dendrites (WT: n = 33 dendritic segments, 226 individual mitochondria, mean length = 6.562 μm 0.437 (SEM), mean occupancy = 84.19% ± 1.44%; Camkk2^-/-^: n = 35 dendritic segments, 199 individual mitochondria, mean length = 11.14 μm ± 0.758 (SEM), mean occupancy = 95.03% ± 1.73%, length increase = 69.77%, occupancy increase = 12.88%; Camkk2^-/-^ + Mtfr1l^S2D^: n = 13 dendritic segments, 110 individual mitochondria, mean length = 5.419 μm ± 0.402 (SEM), mean occupancy = 81.83% ± 2.71%). Scale bar, 5 μm.

These results demonstrate that, in hippocampal neurons, Mtfr1l is phosphorylated by AMPK in an activity- and Camkk2-dependent manner.

Next, we tested whether Camkk2-dependent Mtfr1l phosphorylation by AMPK is required for the compartmentalized mitochondrial morphology characterizing CA1 PNs *in vivo*. Previous work demonstrated that AMPK phosphorylates two conserved serine residues on Mtfr1l (positions S103 and S238), and that a phosphomimetic form of Mtfr1l^S103D/238D^ (hereafter referred to as Mtfr1l^S2D^) is sufficient to rescue mitochondria morphology in AMPK-null cells ^33^. Therefore, we reasoned that if Camkk2 regulates the dendritic compartmentalization of mitochondrial morphology in CA1 PNs by AMPK-dependent phosphorylation of Mtfr1l, expression of phosphomimetic Mtfr1l^S2D^ should rescue the mitochondrial morphology defects observed in Camkk2-null CA1 PNs *in vivo*. Indeed, we found that expression of Mtfr1l^S2D^ in Camkk2-null CA1 PNs rescues the morphology of dendritic mitochondria both in SR and SO compartments back to levels observed in wild-type mice (**Fig. 6B-J**).

## DISCUSSION

In this study, we identified novel molecular and cellular effectors that enable synaptic activity to shape the compartment-specific morphology of mitochondria in neuronal dendrites. In the apical tufts (SLM) of CA1 pyramidal neurons, mitochondria are elongated, tubular, and fill a high fraction of the dendritic volume; in contrast, the mitochondria found in more proximal dendritic compartments are short and occupy a significantly smaller fraction of dendritic volume. This striking compartmentalization of mitochondria morphology corresponds to dendritic domains receiving different presynaptic inputs, from the entorhinal cortex in SLM and from CA2/3 in SR and SO. Collectively, our results (summarized in **Fig. 7**) demonstrate that the mitochondrial morphology observed in the SR and SO dendritic compartments is shaped by synaptic activity via Camkk2- dependent AMPK phosphorylation of the pro-fission Drp1 receptor Mff ^11, 23^ and phosphorylation of the anti-fusion Mtfr1l ^33^, a protein antagonizing Opa1.

**Figure 7.**
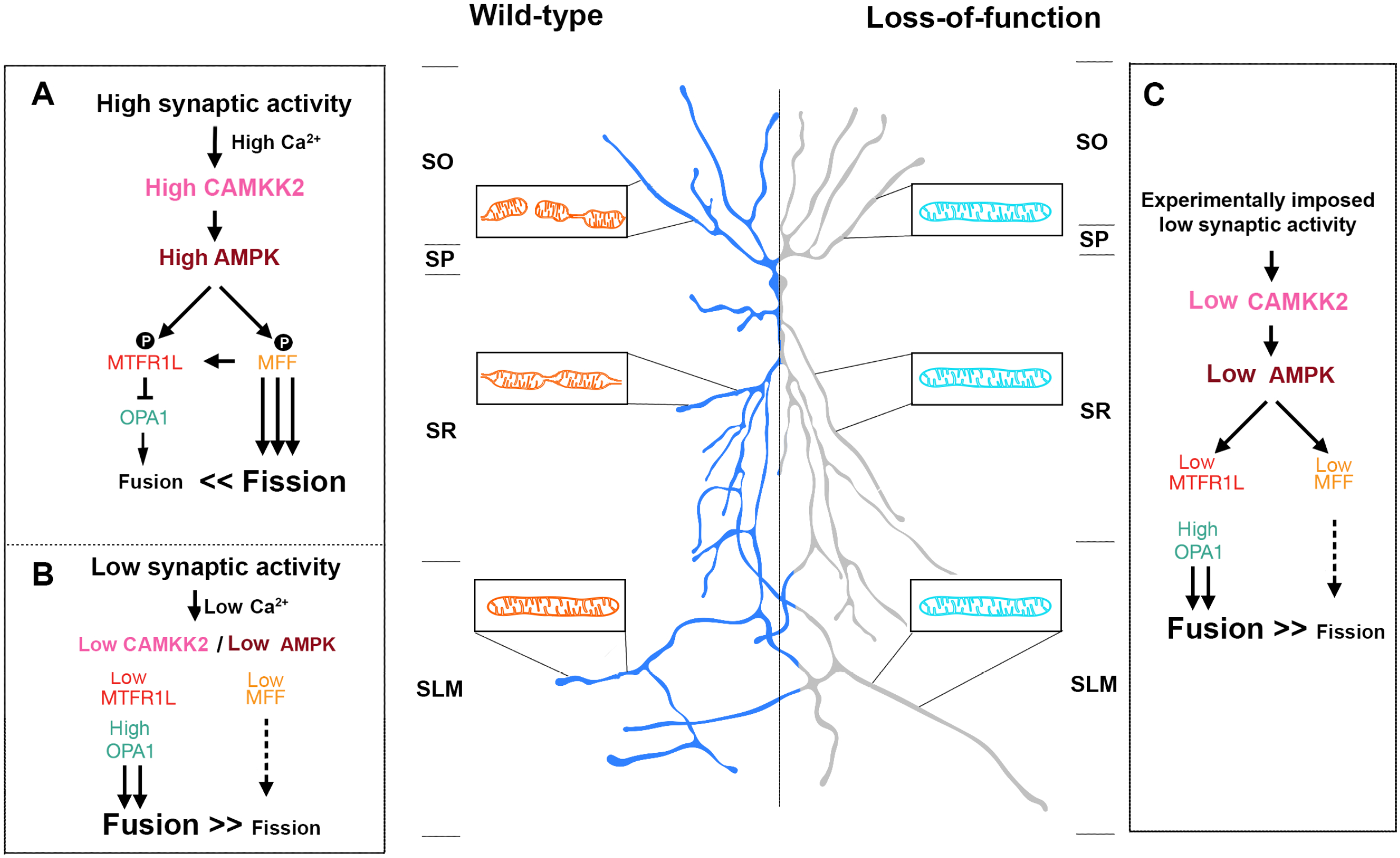
Summary of the main findings. In wild-type mouse CA1 PNs, dendritic mitochondria display a striking degree of compartmentalized morphology, being long and fused in the apical tufts (SLM) with progressive fragmentation and occupancy of a smaller volume of the dendritic segments in SR and SO respectively. We demonstrate using loss-of-function as well as rescue experiments that this compartmentalization of dendritic mitochondria morphology requires (1) neuronal activity (blocked by neuronal hyperpolarization following over-expression of Kir2.1) or by reducing the number of presynaptic inputs from CA3 by ∼50% (shRNA Lphn3) *in vivo*, (2) requires activity-dependent activation of AMPK mediated by Camkk2 and (3) requires the AMPK-dependent phosphorylation of the pro-fission Drp1 receptor Mff and the anti-fusion protein Mtfr1l though its ability to suppress the pro-fusion Opa1 protein. These results demonstrate that mitochondrial fusion dominates over fission in apical tuft dendrites (SLM) and that fission dominates fusion in both SO and SR dendritic compartments. See Discussion for details.

The most parsimonious interpretation of our results is that mitochondrial fusion is dominant in dendrites, and constitutes the ‘default’, as the elongated mitochondria morphology observed in the apical tufts (SLM) of CA1 PNs is present across the entire dendritic arbor in the absence of or under low levels of synaptic activity (**Fig. 7**). In support of this concept, experimentally reducing neuronal activity through expression of Kir2.1 or cell-autonomously reducing by ∼50% the number of CA3 inputs received by individual CA1 PNs (**Fig. 2**), is sufficient to induce tubular and fused mitochondria morphology in the SO and SR dendritic compartments *in vivo* (**Fig. 2**). Therefore, our data suggests that presynaptic activity and cytoplasmic Ca^2+^ dynamics in basal and apical oblique dendrites of CA1 PNs drives the Camkk2 dependent pathway to trigger high levels of AMPK kinase activity and phosphorylation of the anti-fusion effector Mtfr1l (**Fig. 4;** ^33^) and pro-fission Mff (**Fig. 5;** ^11, 23^). In the apical tuft, we hypothesize that lower levels of synaptic activity and/or lower levels of cytoplasmic Ca^2+^ levels due to lower amplitude and/or frequency of Ca^2+^ transients than in SO/SR compartments leads to low levels of Camkk2/AMPK activity and therefore low levels of Mff and Mtfr1l activity (**Fig. 7B**). Our biochemical and rescue experiments (**Fig. 6**) strongly argue in favor of a model whereby Mtfr1l phosphorylation by AMPK is increased by neuronal depolarization (which activates VGCCs and increases intracellular Ca^2+^; ^34^) in a Camkk2-dependent manner.

What could be the functional consequence of this striking compartmentalized mitochondrial morphology in dendrites of CA1 PNs? As mitochondria buffer a significant fraction of the Ca^2+^ released from the ER in dendrites of CA1 PNs ^35^, one potential consequence of the reduced mitochondrial volume in SR/SO dendrites would be that mitochondria in these compartments have a reduced capacity to buffer Ca^2+^, therefore representing a positive-feedback loop allowing more cytoplasmic Ca^2+^ to accumulate in these compartments upon synaptic activity, further increasing Camkk2/AMPK activity. In the apical tufts of the same neurons, lower synaptic activity (spine density is ∼40% lower in SLM than in SO/SR dendrites ^28, 36^), might trigger lower amplitude/frequency of Ca^2+^ transients and therefore lower levels of Camkk2/AMPK activity allowing mitochondrial fusion to dominate over Mff-dependent fission.

The novel molecular effectors identified in the present study will enable future investigations of the functional impact of this striking degree of compartmentalization of mitochondrial morphology on dendritic integration properties, synaptic plasticity, and circuit properties of CA1 PNs *in vivo* ^35^.

## Author Contributions

Conceptualization: FP, TL. Formal analysis: DV, SH, BO, EZ, NP, VH, DZ, KG, FP, TL. Funding acquisition: FP, TL. Investigation: DV, SH, BO, AM, EZ, NP, VH, DZ, KG, EB, FP, TL. Methodology: DV, SH, BO, EZ, KG, EB, FP, TL. Project administration: FP, TL. Resources: EB, FP, TL. Supervision: FP, TL. Visualization: DV, SH, BO, FP, TL. Writing – original draft: DV, SH, FP, TL. Writing – review & editing: DV, SH, BO, EB, FP, TL.

## Acknowledgements

We thank past and present members of the Polleux and Lewis labs for feedback and discussion along the way. We thank Nelson Spruston (HHMI-Janelia) for sharing the CA1 serial EM dataset generated while EB was in his laboratory. We thank Grahame Hardie (Univ. Dundee- UK) and Julien Prudent (Univ. Cambridge- UK) for generously sharing the phospho-specific MTFR1L antibody. We thank Patrycja Szybowska, Joshua Weertman, Klaudia Strucinska, Qiaolian Liu and Rhythm Sharma for technical help. This research was supported by grants NIGMS R35GM137921 (TL), NIGMS P20GM103636-06 sub-project 2 (TL), Presbyterian Health Foundation (TL), NINDS R35 NS127232 (FP) and NINDS NS107483 (FP).

## METHODS

### Animals

All experiments involving mice were done according to protocols approved by the Institutional Animal Care and Use Committee (IACUC) at Columbia University, Oklahoma Medical Research Foundation (OMRF) and in accordance with National Institutes of Health guidelines. Animal health and welfare were supervised by a designated veterinarian. All mice were maintained in Columbia University or OMRF animal facilities that comply with all appropriate standards of care, including cage conditions, space per animal, temperature, humidity, food, water, and 12-hour light/dark cycles.

Timed-pregnant hybrid F1 control females were obtained by mating inbred *129/SvJ* females (Charles River) and *C57BL/6J* males in house. Homozygous double conditional knockout (cKO) lines AMPKα1^F/F^ α2^F/F^ ^37^ were provided by Dr Benoit Viollet (INSERM, Institut Cochin- Paris, France) and homozygous *Camkk2*^-/-^ ^38^ were obtained from Dr. Talal Chatila (Harvard Medical School, Boston). Both AMPKα1^F/F^ α2^F/F^ double cKO and *Camkk2*^-/-^ timed-pregnant females were obtained by mating homozygous males with females of the same genotype. CD-1 IGS mice (Strain Code: 022) were purchased from Charles River Laboratories.

### Cell and Tissue Lysis and Western Blotting

Human Embryonic Kidney 293T/17 (HEK293T) cells were purchased from ATCC (CRL-11268). 1x10^5^ HEK cells were resuspended in media (DMEM, Gibco) with penicillin/streptomycin (0.5×; Gibco) and FBS and seeded in 6 well plates (Corning). Transfection with plasmid DNA (1 mg/mL) using jetPRIME® reagent (Polyplus) according to manufacturer protocol was performed 24 hours after seeding. 72 hours following transfection, cells were carefully washed with 1xPBS (Gibco) then collected into RIPA buffer with protease inhibitor cocktail.

Aliquots of the collected samples were separated by SDS-PAGE and then transferred to a polyvinylidene difluoride (PVDF) membrane (Amersham). After transfer, the membrane was washed 3X in Tris Buffer Saline (10 mM Tris-HCl pH 7.4, 150 mM NaCl) with 0.1% of Tween 20 (T-TBS), blocked for 1 hr at room temperature in Odyssey Blocking Buffer (TBS, LI-COR), followed by 4°C overnight incubation with the appropriate primary antibody in the above buffer. The following day, the membrane was washed 3X in T-TBS, incubated at room temperature for 1 hr with IRDye secondary antibodies (LI-COR) at 1:10,000 dilution in Odyssey Blocking Buffer (TBS), followed by 3X T-TBS washes. Visualization was performed by quantitative fluorescence using an Odyssey CLx imager (LI-COR). Signal intensity was quantified using Image Studio software (LI-COR). Primary antibody used for Western blotting was mouse anti-LPHN3 (R&D Systems MAB5916) and rabbit anti-GAPDH (CST 2118). Total protein was assessed using the Revert 700 Total protein stain (LI-COR 926-11010).

### Activity Assay

*Cell Culture and Lysis:* Embryos were harvested at E15.5 and chilled in Hank’s Balanced Salt Solution (HBSS, Thermo Fisher Scientific) while medial portion of the dorsal telencephalon (presumptive hippocampi) were dissected out. Isolated hippocampi were incubated in papain for 15 minutes at 37°C. After three washes of HBSS, hippocampi were manually dissociated with a pipette into Neurobasal (Thermo Fisher Scientific) supplemented with FBS (Gemini Bio-Products), GlutaMAX (Thermo Fisher Scientific), penicillin/streptomycin (Thermo Fisher Scientific), B27 (Fisher Scientific), and N2 (Thermo Fisher Scientific). Dissociated cells were plated into the wells of 6 well plates (Corning) (7.5 x 10^5^ cells/well) that were previously coated in Poly-D-Lysine (Thermo Fisher Scientific) and incubated at 37°C with 5% CO2. Cells were fed every 3-4 days by removing 0.75mL of media and replenished with 1mL of fresh Neurobasal containing GlutaMAX, B27 and N2 only. Cells were brought to 21 days *in vitro* (DIV) and then treated with either DMSO (2.5µM, Santa Cruz Biotechnology), STO609 (2.5µM, Sigma-Aldrich), or Compound 991 (50µM, Selleck Chem) for 2.5 hours. Neurobasal media was then switched out for 5K Tyrode Solution containing TTX (1µM, Hello Bio) to block sodium channels, NMDA antagonist AP5 (10µM, Hello Bio), and AMPA antagonist NBQX (50µM, Hello Bio) for 30 minutes to dampen cell activity. Media was then switched out for either a second incubation in 5K or 40K Tyrode Solution for 15 minutes. Drug treatments were maintained throughout every incubation step. Tyrode Solution was removed, and cells were lysed on ice with 100µl of N-PER (Thermo Fisher Scientific) containing phosphatase inhibitors (Sigma Aldrich), protease inhibitor (Sigma Aldrich), and benzonase (EMD Millipore). Lysates were then denatured in 4x Laemmli Buffer (BioRad) and 2-Mercaptoethanol (BioRad) at 95°C for 5 minutes.

*Western Blotting:* Equal amounts of lysates were loaded onto Mini-Protean TGX (4-20%) SDS-PAGE gels (BioRad). All blots except phospho-MTFR1L were transferred to nitroceullose membranes (BioRad) with a Trans-Blot Turbo and blocked for one hour in Intercept Buffer (Li-COR). Membranes were then incubated in their respective primary antibodies in blocking buffer at 4°C overnight. Blots probing for phospho-MTFR1L we transferred to polyvinylidene difluoride membranes (PVDF, Immobilion) and blocked in 5% fat-free dry milk in TBS-T for 1 hour before primary antibody incubation for four days at 4°C. Membranes were incubated in Li-Cor fluorescence-coupled secondary antibodies for 1 hour at room temperature prior to visualization with a Li-Cor Odyssey Blot Imager.

*Antibodies:* AMPKα (Cell Signaling, #2532S, 1:2000); phospho-AMPKα (Cell Signaling, #2535S, 1:2000); MFF (Proteintech #17090-1-AP, 1:1000); phospho-MFF (Affinity Bioscience #AF2365, 1:2000), MTFR1L (Atlas Antibodies #HPA027124, 1:2000); phospho-MTFR1L (Gift from the Graham Hardie Lab, S103 and S235, 1:500; characterized in ^33^)

### Lentiviral particle production and transduction

Lentiviral particles were generated as described before (https://www.addgene.org/protocols/lentivirus-production/). Briefly, freshly thawed HEK293T cells (ATCC, CRL-3216) were seeded in a 10 cm dish containing DMEM (GIBCO, 12430-054) supplemented with 10% FBS (Thermo Scientific, 16141002), 1% GlutaMax (GIBCO, 35050061), 1% Sodium Pyruvate (GIBCO, 11360070), and 1% Pen / Strep (Life Technologies, 15140-122), and grown at 37°C, 5% CO2 for ∼ 48 hours. Once confluent, cells were detached from the dish by trypsinization (GIBCO, 25200072), split into 15 cm dishes, and grown to ∼80 % confluence. Before transfection, cultures were treated with 25 μM chloroquine diphosphate for 5 h, supplemented in fresh growth media. For one 15 cm dish, the transfection reaction was assembled in two separate tubes, each containing 1.5 ml serum-free DMEM. The plasmid mixture, composed of pMD2.G (Addgene, #12259), psPAX2 (Addgene, #12260), and a designated transfer vector, was added to the first tube, and the transfection reagent Polyethylenimine (PEI) to the second tube. Both halves were mixed and incubated at RT for 20 min, time after which the reaction was added dropwise to the dish. Cultures were incubated with the transfection reaction for 24 h and then replaced by fresh growth media. Media containing lentiviral particles was collected 72 h after transfection and spun down for 5 min at 2000 g to pellet cell debris. The supernatant was filtered through a 0.45 μm pore, aliquoted in 1.5 ml Eppendorf tubes, and centrifuged for 2 h at 16000 g at 4°C. The supernatant was discarded, and dry viral pellets were stored at -80°C until use. For neuronal transduction, viral pellets were resuspended in serum-free Neurobasal medium (GIBCO, 21103049), added to neuronal cultures, and incubated for 5-7 days before sample collection.

### Single Cell Electroporation (SCE) of CA1 PNs

SCEs were performed as previously described in detail^35, 39^. Briefly, DNA plasmid constructs were diluted to 50ng/µL and electroporated into single dorsal hippocampal CA1-area pyramidal neurons, guided by two-photon microscopy at 920 nm excitation. Successfully electroporated cells were imaged 48-72h post SCE.

### In utero electroporation (IUE) targeting CA1 PNs

In utero electroporation targeting the dorsal hippocampus was performed using a triple-electrode setup as previously described ^36, 40, 41^ in E15.5 mouse embryos in order to target the dividing progenitors generating CA1 pyramidal neurons. Briefly, endotoxin-free plasmids were injected into the lateral ventricle (1 μg/μL followed by 5 pulses at 45V (50ms duration, 500ms inter-pulse interval) using two anodes (positively charged) laterally on either side of the head and one cathode (negatively charged) rostrally at a 0° angle to the horizontal plane. Mice were perfused transcardially with ice cold PFA (2-4%) and glutaraldehyde (0.075%) at P21, post-fixed overnight in the same solution at 4C, and then sectioned at 110-125 μm on a vibratome (Leica). Sections were mounted with Fluoromount-G aqueous mounting medium (ThermoFisher Scientific) and imaged on a Nikon Ti- E A1R laser-scanning confocal microscope using a 40x or 60x NA1.49 objective, or a Zeiss LSM 880 confocal microscope controlled by Zeiss Black software. Imaging required two lasers 488nm and 561nm together with Zeiss objectives 20x (1.2NA) with 2x zoom, or 100x oil (1.25NA) with 3x zoom. Neuronal processes were visualized by Z-stacking that was later processed into Maximum Intensity Projection (MIP). MIP 2D images were then used for analysis of mitochondrial length and occupancy using NIS Elements software (Nikon) or Fiji (Image J).

### Primary neuronal culture

Following *in utero* electroporation, at E18.5 embryonic mouse cortices were dissected in Hank’s Balanced Salt Solution (HBSS) supplemented with HEPES (10LmM, pH 7.4), and incubated in HBSS containing papain (Worthington; 14LU/mL) and DNase I (100 μg/mL) for 15 min at 37 °C with a gentle flick between incubation. Samples were washed with HBSS three times and dissociated by pipetting on the fourth wash. Cells were counted using Countess™ (Invitrogen) and cell suspension was plated on poly-D-lysine (1Lmg/mL, Sigma)-coated glass bottom dishes (MatTek) or poly-D-lysine/laminin coated coverslips (BD bioscience) in Neurobasal media (Gibco) containing FBS (2.5%) (Sigma), B27 (1 X) (Gibco), and Glutamax (1 X) (Gibco). After 7 days, media was changed with supplemented Neurobasal media without FBS, and 1/3 of the media replaced every 3 days.

### Fixation for primary neuron culture

Culture dishes were fixed with 2% PFA (PFA Alfa Aesar)/0.075% GA (Electron Microscopy Science, EMS) in 1x PBS (Sigma) for 10 minutes. Dishes were washed three times following fixation with 1x PBS (Sigma) for 10 minutes.

### Immunohistochemistry

Following fixation and washing, brains were embedded in 3% low melt agarose (RPI, A20070) in 1x PBS. Brains in agarose cubes were sectioned using a vibratome (Leica VT1200) to 120 μm thick sections. Sections/coverslips were then incubated with primary antibodies (chicken anti-GFP Aves Lab 1:4000, rabbit anti-dsRed Abcam 1:4000) that were diluted in the Blocking buffer (1%BSA, 0.2%TritonX-100, 5%NGS in PBS) at 4 °C for 48h. Subsequently sections were washed 6 times for 10 min in PBS and incubated with secondary antibodies (Alexa conjugated goat anti-chicken488 and goat anti-rabbit568 1:4000) at 4 °C for 48h. The excess of secondary antibodies was removed by six, 10min washes in 1xPBS. Sections were then mounted on slides and coverslipped with Aqua PolyMount (PolyMount Sciences, Inc.) and kept at 4 °C.

### Analysis of spine density and mitochondrial morphology

Dendritic spine and mitochondrial morphology quantification was carried out by first identifying 2° or 3° dendritic segments in the basal, apical oblique, or apical tuft dendritic compartments using the cell fill channel only, blinding the experimenter to mitochondrial morphology and density, removing potential bias during segment selection. Individual dendritic segments were then measured in their length manually using Fiji or NISElements software (ImageJ; NIH or Nikon), with each individual mitochondria length in each dendritic segment subsequently measured manually. The same was done for dendritic spines in a select number of segments. In order to calculate mitochondrial occupancy, the sum of an individual dendritic segment’s mitochondria is divided by the total length of the dendritic segment itself. Spine density similarly is calculated by dividing the total number of dendritic spines by the total length of the dendritic segment.

### Plasmids

pCAG:Cre (Addgene: Cat#13775), pEF1α:flex-tdTomato (Addgene: Cat#28306), pCAG:mito-YFP (Addgene; Cat# 168508) ^42^, pCAG:mito-dsRed ^43^, pCAG:tdTomato ^42^, pCAG:mGreenLantern (Addgene: Cat#164469), pCAG:mtYFP-P2A-tdTomato ^42^, pCAG:Kir2.1-T2A-tdTomato was a gift from Massimo Scanziani (Addgene plasmid #60598), pLKO.1:shMTFR1L (GAGTGGAGTGTATCTGCTTAAGGGG) ^33^, pCAG:MTFR1L^WT^ ^33^, pCAG:MTFR1L^S103D/S238D^ ^33^, pLKO.1:shMFF (CCGGGATCGTGGTTACAGGAAATAA – TRCN0000174665) ^7^, pLKO.1:shOPA1 (CCGACACAAAGGAAACTATTT – TRCN0000091111), pLKO.1:shNT (CGTTAATCGCGTATAATACGCGTAT) ^33^. pCMV6 Lphn3 (MR212027), pGFP-C-shLenti scrambled negative control (TR30021), and pGFP-C-shLenti Lphn3 shRNA-D (GTATGTTGGCTTCGCCTTGACACCTACTT, custom) were purchased from Origene.

### In vivo two-photon calcium imaging

We used the same imaging system as described previously ^35, 36^. All images were acquired using a Nikon 40 x NIR water-immersion objective (0.8 NA, 3.5 mm WD) in distilled milliQ water. For excitation, we used a Chameleon, Ultra II (Coherent) laser tuned to 920 nm. We continuously acquired red (tdTomato) and green (GCaMP6f) channels separated by an emission cube set (green: HQ525/70 m-2p; red: HQ607/45 m-2p; 575dcxr, Chroma Technology) at 512 x 512 pixels covering 330 μm x 330 μm at 30 Hz with photomultiplier tubes (green GCaMPF6f fluorescence, GaAsP PMT, Hamamatsu Model 7422-40; red tdTomato fluorescence, GaAsP PMT Hamamatsu).

### Reagents

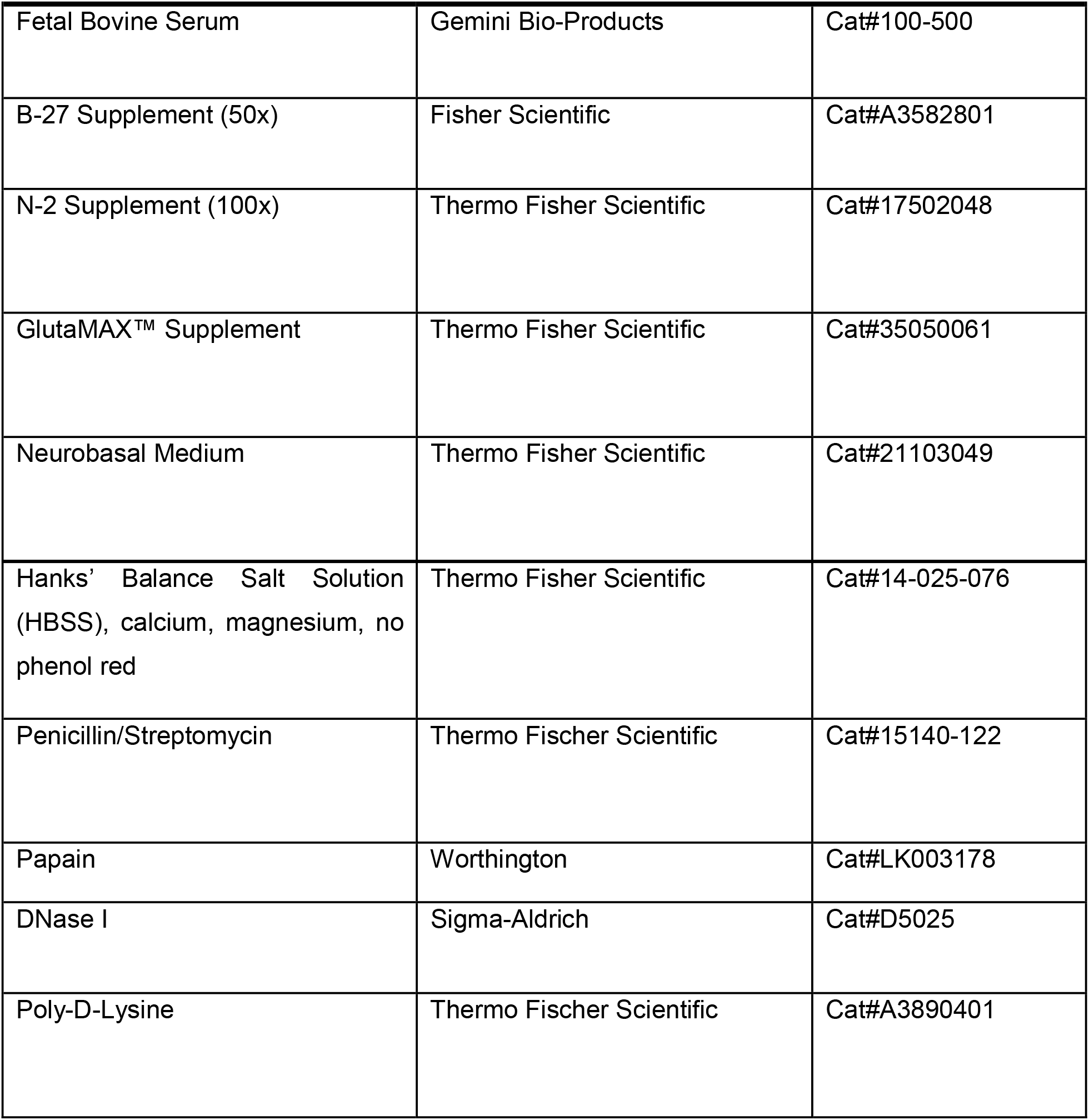
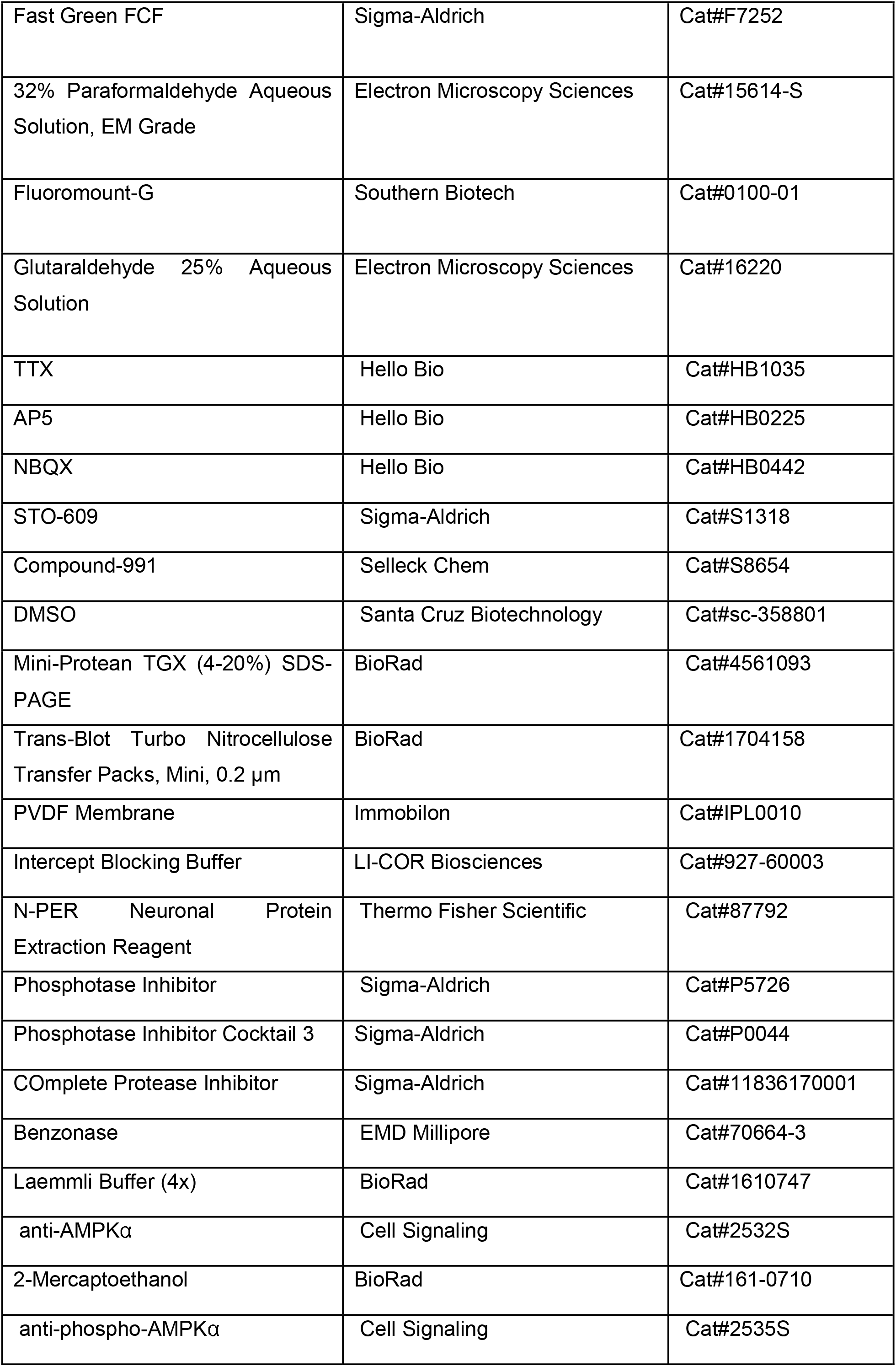
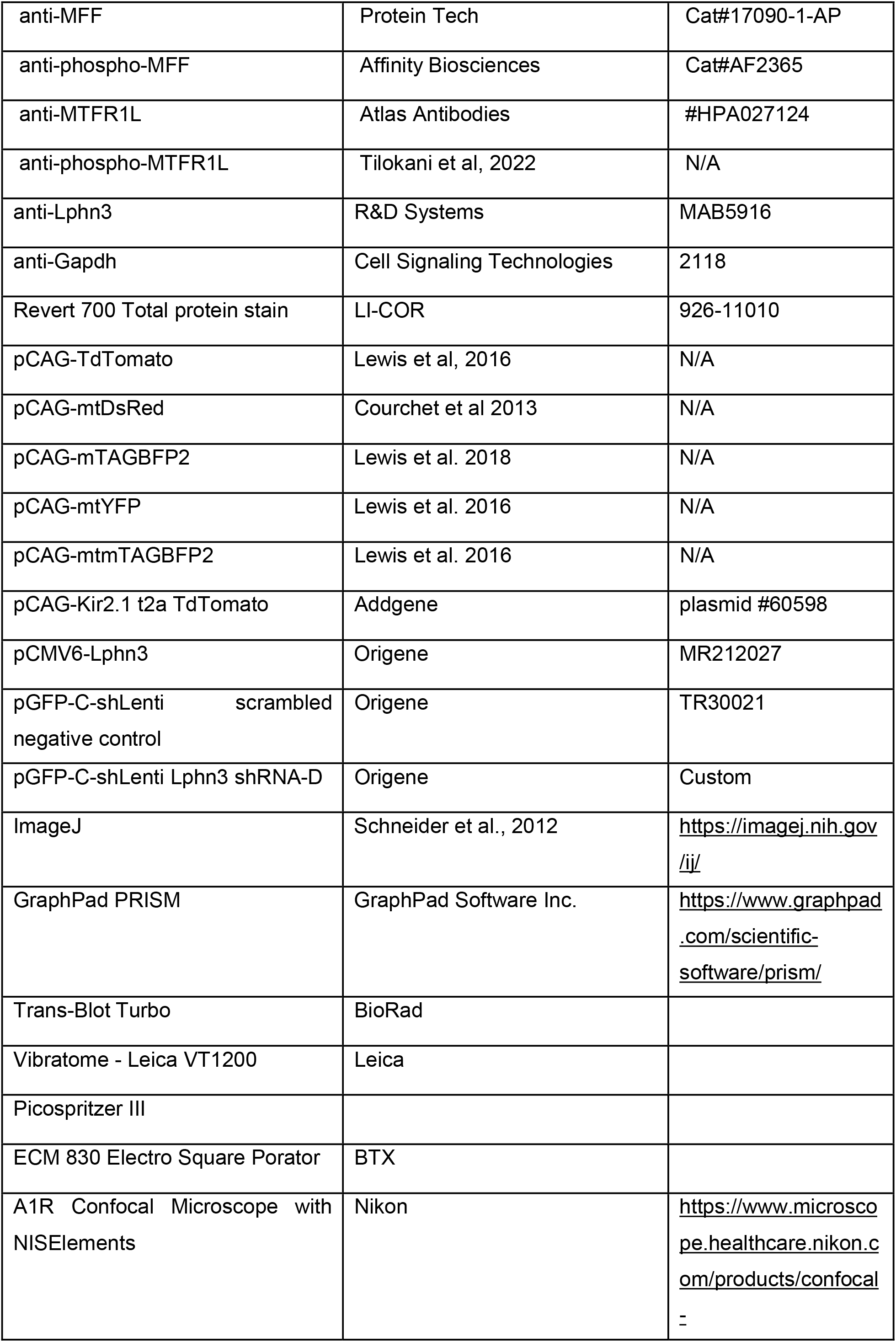
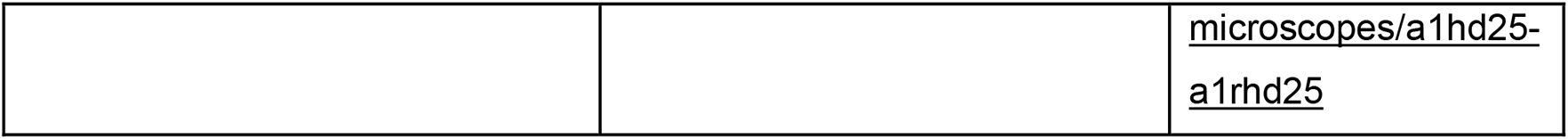

