## Supplemental Figures for "Activity-dependent subcellular compartmentalization of dendritic mitochondria structure in CA1 pyramidal neurons"

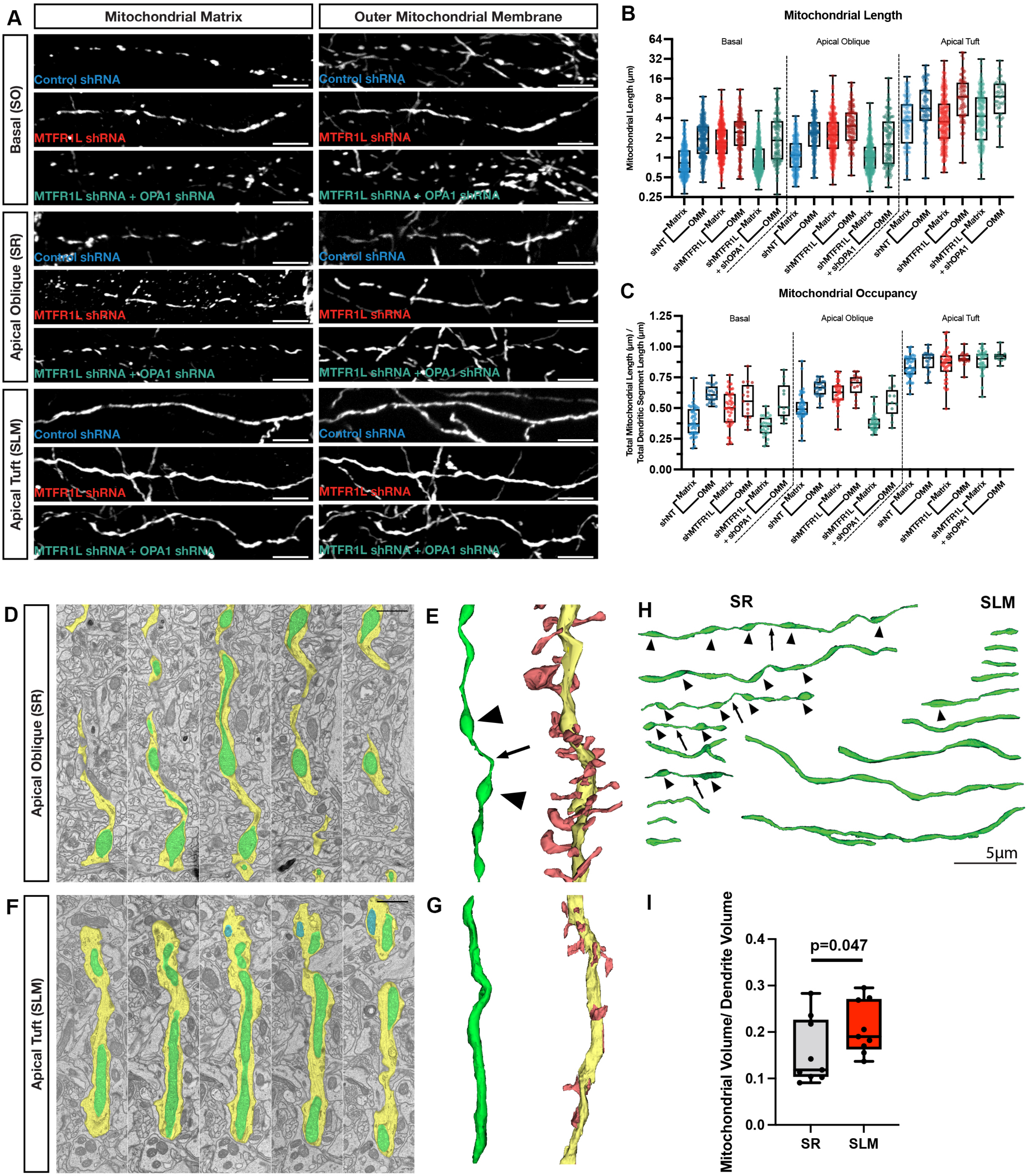

A
Mitochondrial Matrix
Outer Mitochondrial Membrane
Basal (SO)
Apical Oblique (SR)
Apical Tuft (SLM)
Control shRNA
MTFR1L shRNA
MTFR1L shRNA + OPA1 shRNA
B
Mitochondrial Length
Mitochondrial Length (µm)
Basal
Apical Oblique
Apical Tuft
shNT
shMTFR1L
shMTFR1L + shOPA1
Matrix
OMM
C
Mitochondrial Occupancy
Total Mitochondrial Length (µm) / Total Dendritic Segment Length (µm)
D
E
F
G
H
SR
SLM
5µm
I
Mitochondrial Volume/ Dendrite Volume
p=0.047
SR
SLM

**Supplemental Figure 1: Compartmentalized mitochondrial morphology visualized using fluorescent OMM and matrix markers as well as 3D serial electron microscopy (3D-SEM).**
(**A-C**) High magnification representative images of mitochondrial morphology within isolated secondary or tertiary hippocampal CA1 basal, apical oblique, and apical tuft dendrites in which a mitochondrial matrix-targeted fluorescent protein (mt-mTAGBFP2 or mt-YFP), an outer mitochondrial matrix (OMM)-targeted fluorescent protein (mt-ActAmCherry-HA), and cell fill (mGreenLantern or mTAGBFP2—not pictured) were *in utero* electroporated along with either a control shRNA (shNT), *Mtfr1l* shRNA (shMtfr1l), or *Mtfr1l* and *Opa1* shRNA (shMtfr1l + shOpa1) (A). Quantification of mitochondrial length and occupancy in the basal, apical oblique, and apical tuft dendritic compartments reveals a significant increase in mitochondrial matrix length (B) and occupancy (C) in basal and apical oblique dendrites when knocking down Mtfr1l, which is rescued when also knocking down Opa1 (**Fig. S5E-F**). Note the significant increase in OMM length (B) and occupancy (C) in all conditions in all dendritic compartments when comparing against mitochondrial matrix length and occupancy. Nevertheless, knocking down shMtfr1l continues to moderately, but significantly, increase the length (B) and occupancy (C) of the OMM when knocking down Mtfr1l, which is rescued when simultaneously knocking down Opa1. There are no significant differences between shNT, shMtfr1l, or shMtfr1l + Opa1 matrix lengths (B) or occupancies (C), nor between OMM lengths or occupancies. Scale bar: 5 μm.
(**D-G**) Serial transmission electron microscopy images from the apical oblique dendrites (SR, panel D) and the apical tuft (SLM, panel F) where the cytoplasm is annotated in yellow and mitochondria in green. Three dimensional reconstructions of the dendritic segments shown in D and F are shown in panels E and G where the mitochondria are highlighted in green (left) and the dendritic shaft in yellow with dendritic spines in red. In panel E, arrowheads indicate matrix bulges along the continuous mitochondria and the arrow points to a constriction separating two matrix bulges.
(**H**) Multiple examples of 3D reconstructed, individual dendritic mitochondria displaying various morphologies and length from SR (left) and SLM (right). Note the presence of frequent matrix bulges (arrowheads) and thin constrictions (arrows) in mitochondria from SR dendritic segments that are rarely observed in SLM dendritic segments.
(**I**) Quantification of mitochondrial occupancy defined by volume of dendritic segments reconstructed occupied divided by volume occupied by mitochondria in SR and SO. * $p=0.047$ according to one-tailed Mann-Whitney non-parametric test ($U=21$). Number of dendritic segments and mitochondria reconstructed: SR, $n=76$ individual mitochondria along 9 dendritic segments. SLM, $n=101$ mitochondria along 9 dendritic segments.

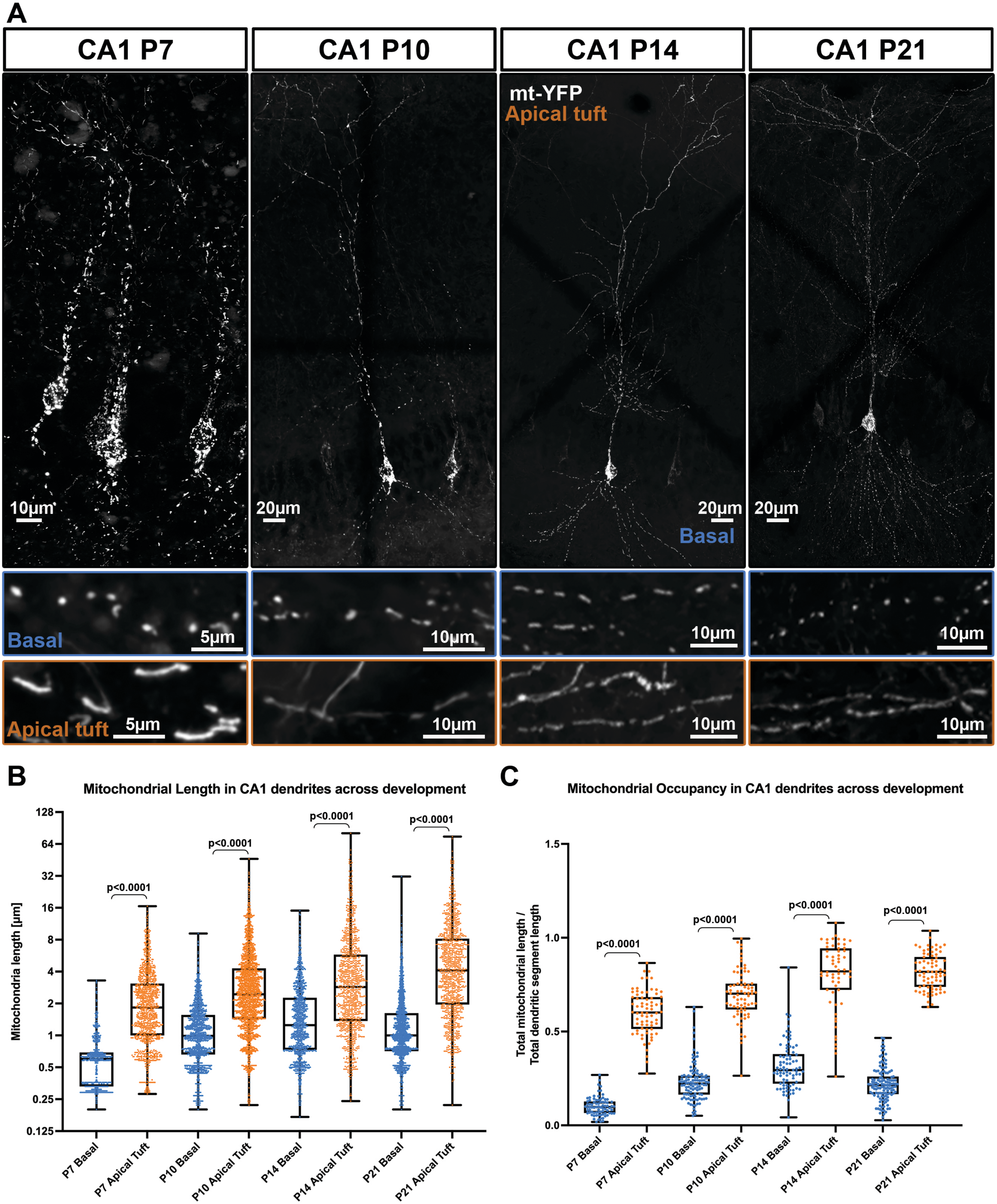
A
CA1 P7
CA1 P10
CA1 P14
CA1 P21
mt-YFP
Apical tuft
Basal
10µm
20µm
20µm
20µm
Basal
5µm
10µm
10µm
10µm
Apical tuft
5µm
10µm
10µm
10µm
B
Mitochondrial Length in CA1 dendrites across development
Mitochondria length [µm]
128
64
32
16
8
4
2
1
0.5
0.25
0.125
p<0.0001
p<0.0001
p<0.0001
p<0.0001
P7 Basal
P7 Apical Tuft
P10 Basal
P10 Apical Tuft
P14 Basal
P14 Apical Tuft
P21 Basal
P21 Apical Tuft
C
Mitochondrial Occupancy in CA1 dendrites across development
Total mitochondrial length / Total dendritic segment length
1.5
1.0
0.5
0.0
p<0.0001
p<0.0001
p<0.0001
p<0.0001
P7 Basal
P7 Apical Tuft
P10 Basal
P10 Apical Tuft
P14 Basal
P14 Apical Tuft
P21 Basal
P21 Apical Tuft

**Supplemental Figure 2: *In vivo* time course of mitochondrial size and occupancy in CA1 pyramidal neurons.**

(**A**) Top: Low magnification representative images of mitochondrial morphologies across CA1 pyramidal neurons at the indicated stages of development following IUE of pCAG mtYFP-P2A-tdTomato at E15.5. Bottom: Higher magnification boxes of mitochondria morphologies from basal (blue) or apical tuft (orange) dendrites at the developmental stage indicated.

(**B-C**) Quantification of mitochondrial length (B) and occupancy (C) in the basal and apical tufts at the indicated timepoints showing clear, distinct regulation of mitochondrial size and occupancy in the basal dendrites and apical tuft dendrites early in development that is maintained into adulthood. $P7_{basal}$ = 79 segments, 759 mitochondria, mean length = 0.63µm ± 0.05µm (SEM), mean occupancy = 10.2% ± 0.5% (SEM); $P7_{apical\ tuft}$ = 72 segments, 1119 mitochondria, mean length = 2.45µm ±0.06µm, mean occupancy = 60.1% ± 1.4%; $P10_{basal}$ = 108 segments, 1622 mitochondria, mean length = 1.23µm ±0.02µm, mean occupancy = 23% ± 0.9%; $P10_{apical\ tuft}$ = 73 segments, 2141 mitochondria, mean length = 3.56µm ± 0.08µm, mean occupancy = 69.5% ± 1.5%; $P14_{basal}$ = 82 segments, 1265 mitochondria, mean length = 1.79µm ± 0.05µm, mean occupancy = 31.6% ± 1.4%; $P14_{apical\ tuft}$ = 59 segments, 1022 mitochondria, mean length = 5.1µm ±0.22µm, mean occupancy = 79.6% ± 2.4%; $P21_{basal}$ = 127 segments, 2154 mitochondria, mean length = 1.42µm ±0.03µm, mean occupancy = 21.8% ± 0.7%; $P21_{apical\ tuft}$ = 75 segments, 1036 mitochondria mean length = 6.48µm ±0.23µm, mean occupancy = 81.9% ± 1.1%. P21 data is the same as the $control_{tdTomato}$ set used in Figure 2. p values are displayed within the figure. Data are shown as individual points on min to max box plots with $25^{th}$, $50^{th}$ and $75^{th}$ percentiles.

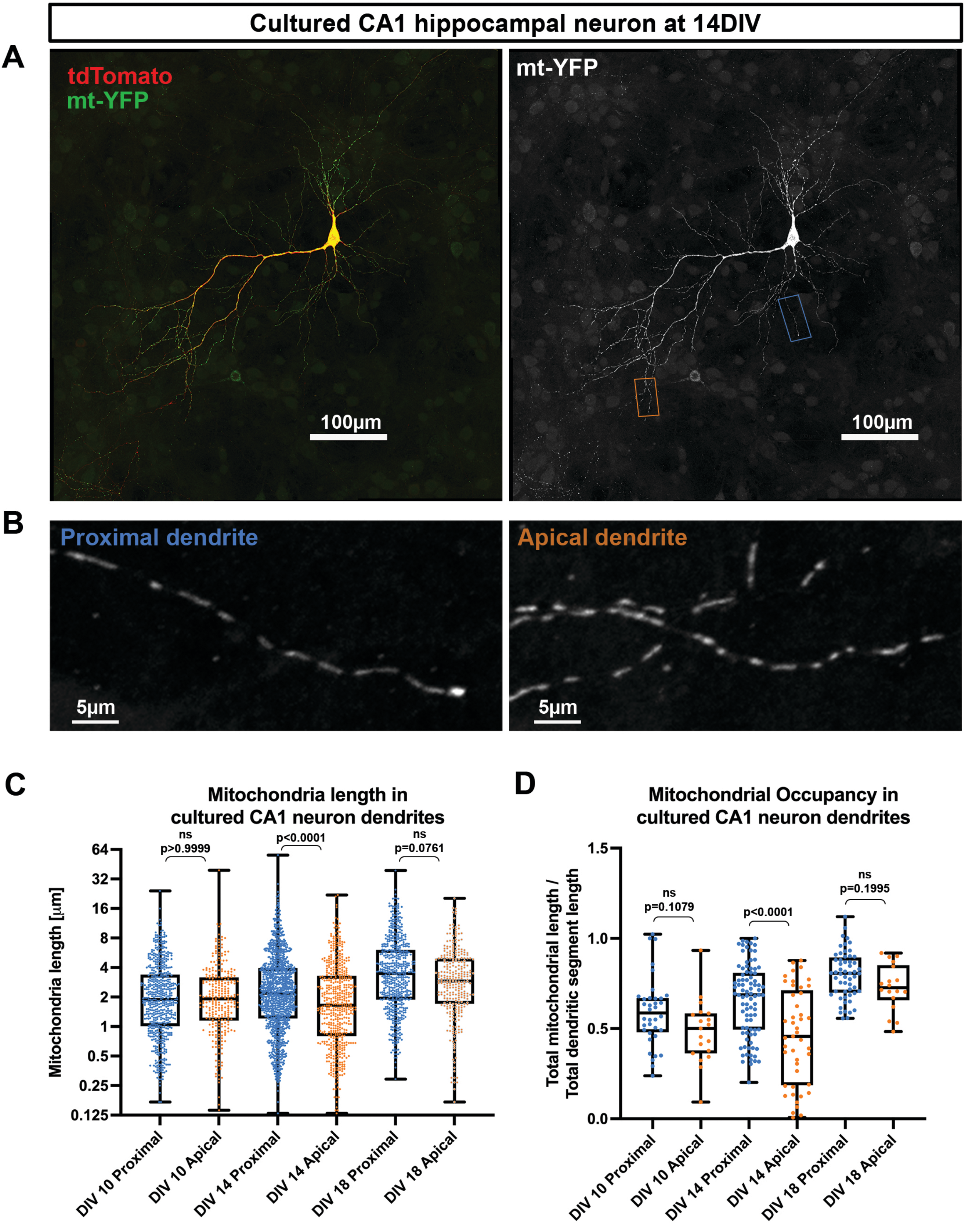
Cultured CA1 hippocampal neuron at 14DIV
A
tdTomato
mt-YFP
100µm
mt-YFP
100µm
B
Proximal dendrite
5µm
Apical dendrite
5µm
C
Mitochondria length in
cultured CA1 neuron dendrites
Mitochondria length [µm]
64
32
16
8
4
2
1
0.5
0.25
0.125
ns
p>0.9999
p<0.0001
ns
p=0.0761
DIV 10 Proximal
DIV 10 Apical
DIV 14 Proximal
DIV 14 Apical
DIV 18 Proximal
DIV 18 Apical
D
Mitochondrial Occupancy in
cultured CA1 neuron dendrites
Total mitochondrial length /
Total dendritic segment length
1.5
1.0
0.5
0.0
ns
p=0.1079
p<0.0001
ns
p=0.1995
DIV 10 Proximal
DIV 10 Apical
DIV 14 Proximal
DIV 14 Apical
DIV 18 Proximal
DIV 18 Apical

**Supplemental Figure 3: Dendritic mitochondria morphology observed in CA1 pyramidal neurons *in vivo* are not conserved in cultured CA1 PNs *in vitro*.**

(**A**) Low magnification representative images of mitochondrial morphology across a hippocampal neuron in culture following IUE of pCAG mtYFP-P2A-tdTomato at E15.5, followed by culture at E18.5. (**B**) Higher magnification of the indicated boxes in A.

(**C-D**) Quantification of mitochondrial length (C) and occupancy (D) in the proximal and apical dendrites at the indicated timepoints showing that in culture the distinct mitochondrial populations observed *in vivo* are absent. $10DIV_{proximal}$ = 36 segments, 580 mitochondria, mean length = 2.64μm ±0.1μm (SEM), mean occupancy = 59% ± 3.2% (SEM); $10DIV_{apical}$ = 19 segments, 275 mitochondria, mean length = 2.58μm ± 0.18μm, mean occupancy = 48.4% ± 4.1%; $14DIV_{proximal}$ = 88 segments, 1199 mitochondria, mean length = 3.2μm ± 0.1μm, mean occupancy = 65.7% ±2.1%; $14DIV_{apical}$ = 45 segments, 527 mitochondria, mean length = 2.56μm ±0.12μm, mean occupancy = 45.5% ± 3.9%; $18DIV_{proximal}$ = 56 segments, 572 mitochondria, mean length = 4.74μm ± 0.18μm, mean occupancy = 79.7% ± 1.7%; 18 $DIV_{apical}$ = 18 segments, 300 mitochondria mean length = 3.7μm ± 0.17μm, mean occupancy = 72.9% ± 3%. p values are shown in the figure. Data are shown as individual points on min to max box plots with $25^{th}$, $50^{th}$ and $75^{th}$ percentiles.

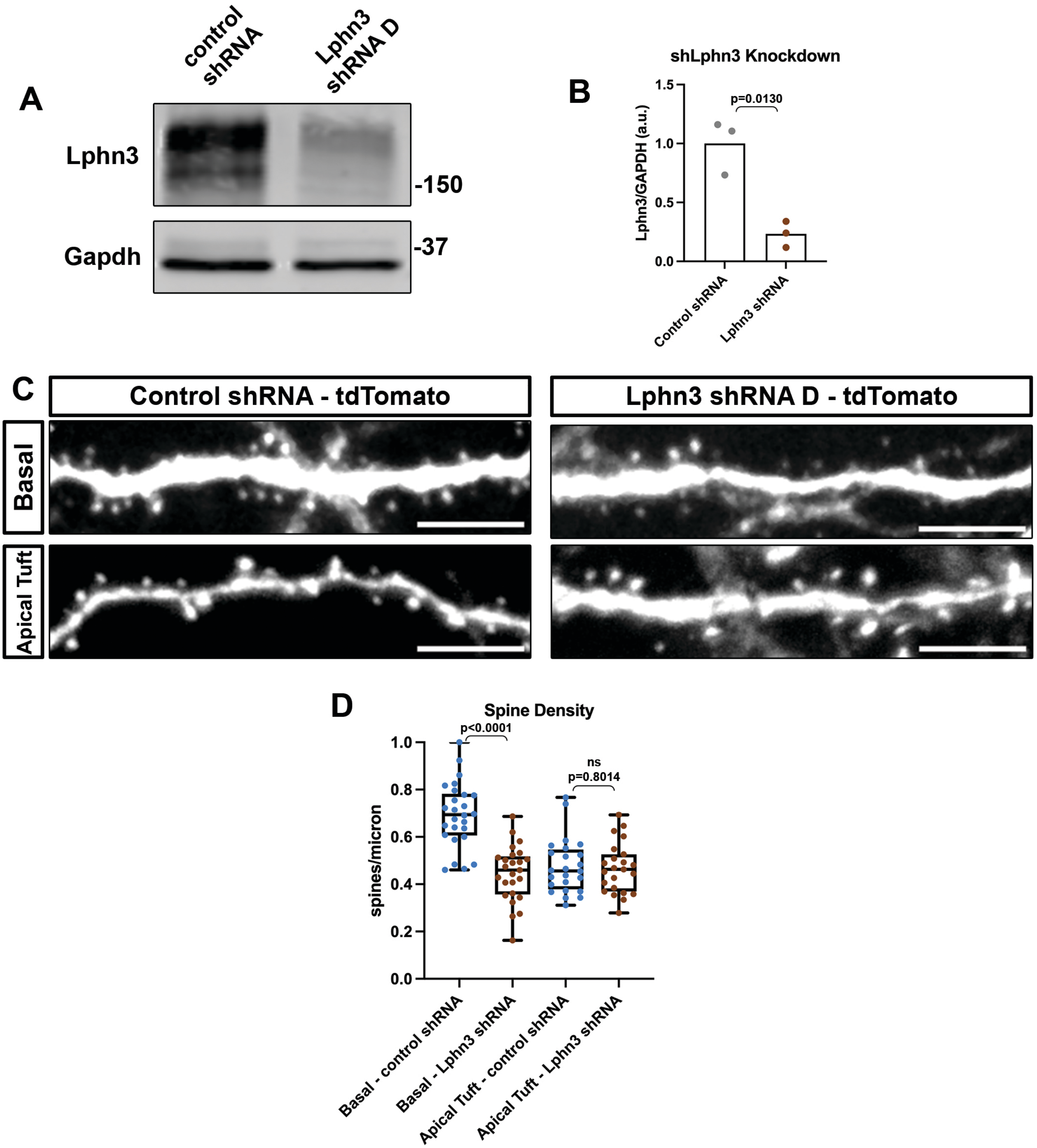
A
control shRNA
Lphn3 shRNA D
Lphn3
Gapdh
-150
-37
B
shLphn3 Knockdown
Lphn3/GAPDH (a.u.)
p=0.0130
1.5
1.0
0.5
0.0
Control shRNA
Lphn3 shRNA
C
Control shRNA - tdTomato
Lphn3 shRNA D - tdTomato
Basal
Apical Tuft
D
Spine Density
spines/micron
p<0.0001
ns
p=0.8014
1.0
0.8
0.6
0.4
0.2
0.0
Basal - control shRNA
Basal - Lphn3 shRNA
Apical Tuft - control shRNA
Apical Tuft - Lphn3 shRNA

**Supplemental Figure 4: Validation of Lphn3 shRNA knockdown construct.**

**A**) Western blot using a mouse Lphn3 antibody (R&D research, top panel) on samples from HEK cells that were transfected with mouse Lphn3 cDNA and either a control or *Lphn3* shRNA construct. Gapdh (Cell Signaling Technologies, bottom panel) was used as a loading control **B**) Relative Lphn3 expression following 72hrs knockdown from 3 independent experiments. Paired t test. p value indicated in the figure. **C**) High magnification representative images of P21 CA1 dendrite segments electroporated at E15.5 with tdTomato and either control shRNA (left) or Lphn3 shRNA D (right) from basal dendrites in SO (top) or apical tuft dendrites (bottom). **E**) Quantification of spine density following IUE with control shRNA or Lphn3 shRNA D at P21 demonstrating that Lphn3 knockdown throughout development results in decreased spine density on basal dendrites but not apical tuft dendrites. Control $shRNA_{basal}$ = 26 dendrite segments, mean spine density = 0.69 ±0.03 spines/micron (SEM), Control $_{shRNAapical\ tuft}$ = 24 dendrite segments, mean spine density = 0.48 ±0.02 spines/micron, Lphn3 $shRNA_{basal}$ = 25 dendrite segments, mean spine density = 0.45 ±0.02 spines/micron, Lphn3 $shRNA_{apical\ tuft}$ = 23 dendrite segments, mean spine density = 0.47 ±0.02 spines/micron. p values are shown in the figure. Data are shown as individual points on min to max box plots with $25^{th}$, $50^{th}$ and $75^{th}$ percentiles. Scale bars are 5 microns.

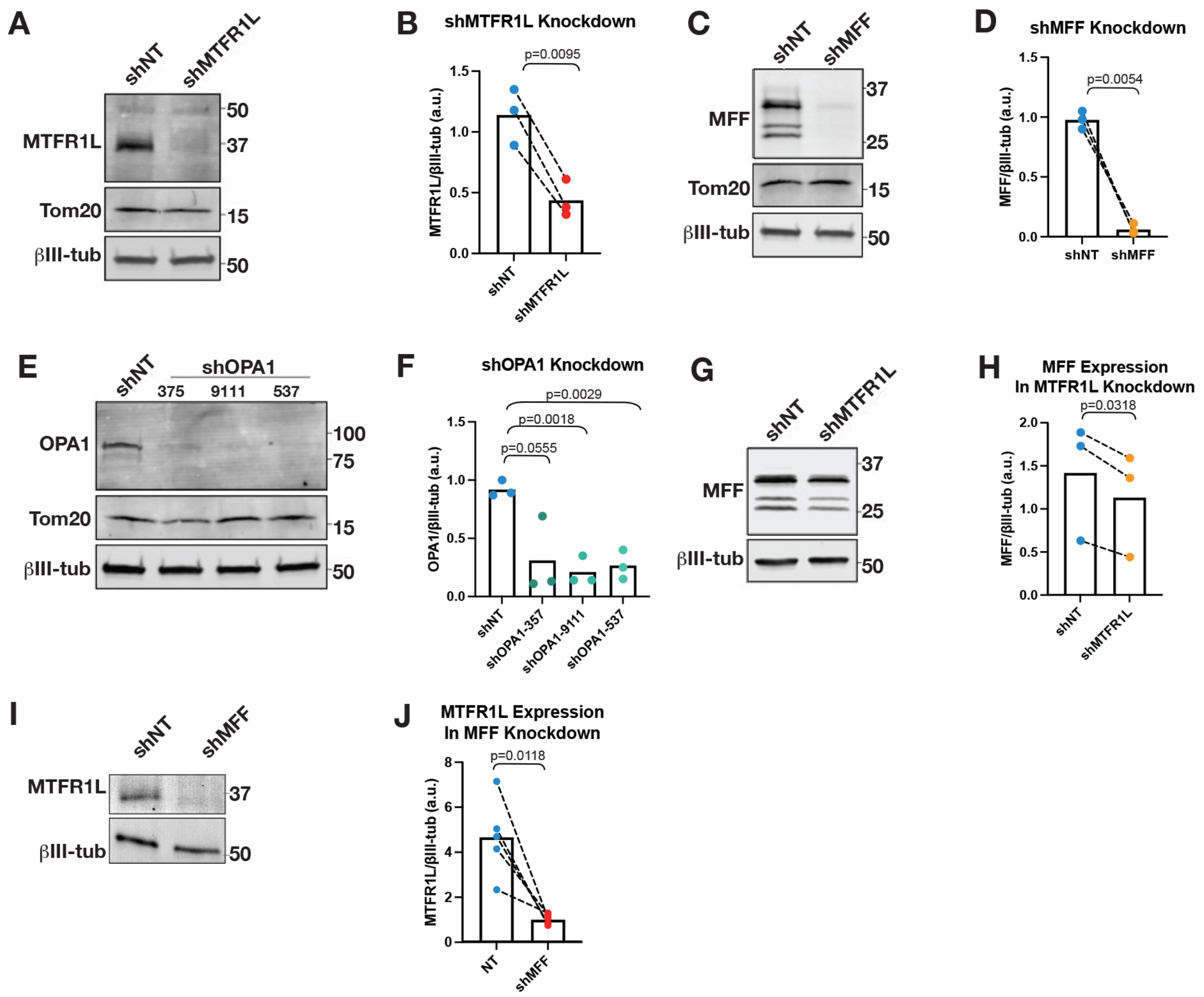
A
shNT
shMTFR1L
MTFR1L
Tom20
βIII-tub
50
37
15
50
B
shMTFR1L Knockdown
p=0.0095
MTFR1L/βIII-tub (a.u.)
1.5
1.0
0.5
0.0
shNT
shMTFR1L
C
shNT
shMFF
MFF
Tom20
βIII-tub
37
25
15
50
D
shMFF Knockdown
p=0.0054
MFF/βIII-tub (a.u.)
1.5
1.0
0.5
0.0
shNT
shMFF
E
shNT
shOPA1
375
9111
537
OPA1
Tom20
βIII-tub
100
75
15
50
F
shOPA1 Knockdown
p=0.0029
p=0.0018
p=0.0555
OPA1/βIII-tub (a.u.)
1.5
1.0
0.5
0.0
shNT
shOPA1-357
shOPA1-9111
shOPA1-537
G
shNT
shMTFR1L
MFF
βIII-tub
37
25
50
H
MFF Expression
In MTFR1L Knockdown
p=0.0318
MFF/βIII-tub (a.u.)
2.0
1.5
1.0
0.5
0.0
shNT
shMTFR1L
I
shNT
shMFF
MTFR1L
βIII-tub
37
50
J
MTFR1L Expression
In MFF Knockdown
p=0.0118
MTFR1L/βIII-tub (a.u.)
8
6
4
2
0
NT
shMFF

**Supplemental Figure 5: Validation of Mtfr1l, Mff, and Opa1 shRNA constructs.**

(**A-B**) Western blot demonstrating shMtfr1l significantly knocks down Mtfr1l (37kDa) compared to control shNT in neuronal cultures. β-Tubulin III (βIII-tub, 50kDa) was stained as a control for neuronal content and Tom20 (15kDa) to control for mitochondrial content and quantification (B) of shMtfr1l knockdown measuring fluorescence intensity of anti-Mtfr1l normalized to anti-βIII-tub.

(**C-D**) Western blot demonstrating shMff significantly knocks down Mff (37-25kDa) compared to control shNT and quantification (D) of shMff knockdown measuring fluorescence intensity of anti-Mff normalized to anti-βIII-tub.

(**E-F**) Western blot demonstrating three separate shOpa1s (357, 9111, 537) significantly knocks down Opa1 (78kDa) compared to control shNT and quantification (F) of shOpa1 knockdowns measuring fluorescence intensity of anti-Opa1 normalized to anti-βIII-tub.

(**G-H**) Western blot probing for Mff in control shRNA and Mtfr1l shRNA treated neurons and quantification (H) of fluorescence intensity of anti-Mff compared to anti-βIII-tub from G.

(**I-J**) Western blot probing for Mtfr1l in control shRNA and Mff shRNA treated neurons and quantification (J) of fluorescence intensity of anti-Mtfr1l compared to anti-βIII-tub from I.
